# A cilia-dependent inflammatory programme links bacterial detection to kidney disease

**DOI:** 10.64898/2026.04.24.720658

**Authors:** Amandine Aka, Joran Martin, Fanny Kopp, Teresa Bulfone, Samuel Chauvin, Camille Cohen, Martine Burtin, E. Wolfgang Kuehn, Anne-Sophie Dangeard, Maryse Moya-Nilges, Nicolas Cagnard, Nicolas Goudin, Meriem Garfa-Traoré, Marceau Quatredeniers, Cong Xu, Laurent Audry, Jost Enninga, Matthieu Rousseau, Cécile Arrieumerlou, Jieqing Fan, Sophie Saunier, Fabiola Terzi, Molly A. Ingersoll, Henri Lichenstein, Amandine Viau, Frank Bienaimé

## Abstract

The primary cilium is a microtubule-based sensory organelle projecting from the plasma membrane of most mammalian cells. Genetic defects in ciliary components cause chronic kidney disease (CKD) characterized by heightened production of inflammatory and fibrogenic mediators by tubular epithelial cells. Yet, whether this reflects a physiological role of the primary cilium remains unknown. Here, we show that primary cilia on kidney tubular cells bind uropathogenic *Escherichia coli* and, in response to bacterial components, initiate a fibro-inflammatory program reminiscent of CKD. Integrating single-cell transcriptomics with conditional mouse models, we observed that epithelial cilia orchestrate a similar fibro-inflammatory response in the absence of infection during CKD. This convergence reveals a shared cilia-dependent signalling axis governing both host–pathogen responses and CKD progression. Mechanistically, cilia ablation selectively impairs tubular responses to ADP-heptose, a pathogen-associated molecular pattern that activates NF-κB via the cytosolic innate immune receptor ALPK1. In human kidney organoids, ADP-heptose induces robust fibro-inflammation, and genetic or pharmacological inhibition of ALPK1 attenuates this response in a rodent CKD model. Together, these findings identify primary cilia as central orchestrators of a fibro-inflammatory program linking pathogen detection to kidney disease progression.

## INTRODUCTION

Rooted in the centrosome, primary cilia are solitary protrusions of the plasma membrane present on most mammalian cells. These microtubule-based organelles serve as sensory structures that detect environmental cues and as signalling platforms regulating essential cellular processes, including growth, autophagy, and metabolism^1^. Primary cilia are highly compartmentalized, with the transition zone at their base acting as a gatekeeper for protein trafficking in and out of the cilia. Distal to the transition zone lies the inversin compartment, which ends before the ciliary tip. This region is defined by a fibrillary protein complex that includes INVS, ANKS6, NEK8, and NPHP3^2^.

Mutations affecting ciliary components cause a group of multisystemic disorders collectively known as ciliopathies^1^. The kidney is particularly susceptible, with renal ciliopathies typically presenting as severe interstitial fibrosis and/or cystic dilation of renal tubules, and usually progressing to kidney failure^3,4^. A hallmark of these disorders is the aberrant secretion of inflammatory and fibrotic cytokines by tubular epithelial cells, which promotes immune cell recruitment, myofibroblast activation, and tissue scarring^5,6^.

In kidney tubule epithelial cells, primary cilia project into the lumen, where they are subjected to deflection by urinary flow. This anatomical position has long supported the hypothesis that cilia function as mechanosensors of fluid flow^7^. *In vitro* studies showed that primary cilia are required for the inhibition of mTORC1 and the activation of autophagy in response to fluid shear stress, leading to cell size reduction^8,9^. However, none of these signalling events have been directly linked to the fibro-inflammatory responses commonly observed in renal ciliopathies.

Fibro-inflammation is a fundamental response to tissue injury implicated in both acute and chronic kidney diseases (CKD). Notably, CKD mechanisms intersect with those governing pathogen defence. Specifically, innate immune pathways that recognize pathogen-associated molecular patterns (PAMPs) have been implicated in CKD progression^10^. Genome-wide association studies reveal that important genetic variants in *APOL1*^11^ and *UMOD*^12^ predisposing to CKD also confer protection against trypanosomes and urinary tract infections (UTI)^11,12^, respectively, supporting evolutionary positive selection. Therefore, we hypothesized that if primary cilia play a role in kidney fibro-inflammation, this function would be unmasked in the context of infection. We considered UTI, caused by uropathogenic *E. coli* (UPEC), as the most prevalent kidney infection. Infection of the kidney most commonly results from the ascent of UPEC from an infected bladder into the lumen of kidney tubules, where cilia protrude^13^. In this work we explored the link between primary cilia, urinary tract infection an CKD progression.

## RESULTS

### Kidney primary cilia bind uropathogenic *E. coli*

To explore whether primary cilia can directly interact with uropathogenic bacteria, we assessed UPEC adhesion to cilia in different tubular cell lines derived from dog, mouse, or human (*i.e.*, MDCK, mIMCD3 and HK2, respectively). Cells were infected with UPEC (UTI89 strain) for two hours, and bacterial attachment to cilia was quantified. UPEC displayed a strong preference for the surface of primary cilia in each of the tested renal tubular cell lines (**Fig. 1A, B**). By contrast, UPEC did not significantly adhere to the primary cilia of human retina-derived epithelial cells (**Fig. 1B**).

**Figure 1.**
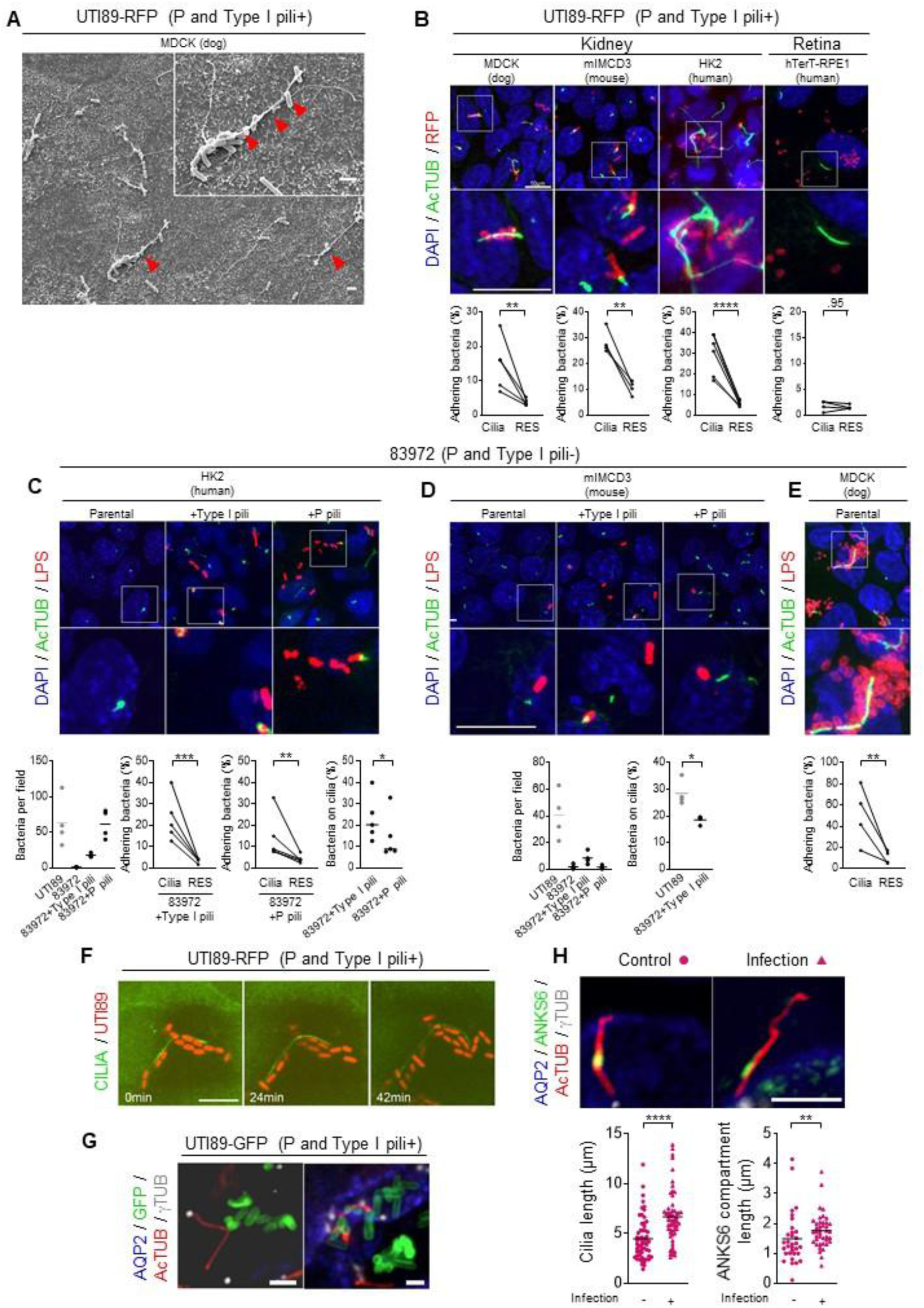
Kidney primary cilia bind uropathogenic *E. coli*. **(A)** Scanning electron microscopy of dog kidney epithelial cells (MDCK), 2 hours after uropathogenic *E. coli* (UPEC, UTI89-RFP strain) infection showing bacteria bound to a primary cilium (arrowheads). Scale bar: 2 µm. **(B)** Representative immunofluorescence images and quantification of the interaction between primary cilia axoneme (AcTUB+, green) and UPEC strain UTI89 (RFP+, red) in kidney cells from dog (MDCK), mouse (mIMCD3), and human (HK2) or retina cells from human (hTerT-RPE1). Scale bar: 10µm. UPEC localisation was recorded as on the primary cilium or at a random equivalent surface for each experiment (RES; cilia area flipped horizontally and vertically, see methods). Each pair of linked dots indicates an independent experiment. **(C)** Representative immunofluorescence images and quantification of the interaction between human kidney cells (HK2) primary cilia (AcTUB+, green) and UPEC strain 83972 (LPS+, red), complemented with type 1 or P-pili and quantification of the absolute number of bacteria per field (left), UPEC localisation on the primary cilium or at a RES in 83972 expressing type 1 or P-pili (middle) and comparison of the percentage of bacteria at cilia between the two lines, Scale bar: 10µm. (**D**) Representative immunofluorescences and quantification of the interaction between mouse kidney cells (mIMCD3) primary cilia (AcTUB+, green) and UPEC strain 83972 (LPS+, red), complemented with type 1 or P-pili and quantification of the absolute number of bacteria per field and the percentage of bacteria located at cilia (left), UTI89. Results obtained with UTI89 are shown as comparator. (**E**) Representative immunofluorescences and quantification of the interaction between dog kidney cells (MDCK) primary cilia (AcTUB+, green) and UPEC strain parental 83972 (LPS+, red). **(F)** Time-lapse imaging of cilia (STRADB+, green) and UPEC UTI89 (red) in MDCK kidney cells (T0=2 hours post infection; see also **Extended Data Movies 1-2**). Scale bar: 10 µm. **(G)** Representative immunofluorescence images demonstrating interactions between primary cilia axoneme (AcTUB+, red) stemming from the γTUB+ basal body (grey), and UPEC strain UTI89 (green) in the collecting duct (AQP2+, blue) of C3H/HeN mice, 5 days after intravesical instillation of UPEC. Scale bars: 3 µm. See also **Extended Data Movies 3-4**. **(H)** Representative images and quantification of primary cilia axoneme length (AcTUB+, red), stemming from the γTUB+ basal body (grey) and ANKS6 (green) compartment length in collecting ducts (AQP2+) in control and infected mice five days after bladder instillation of UTI89 (uropathogenic *E. coli*). Scale bar: 5 µm. Each dot represents a cilium (n=3 mice per group). **(B-E, H)** Bars indicate mean. Ratio paired *t* test (B-C, E), or Mann-Whitney t-test (D, F): *P<0.05, **P<0.01, ***P<0.001, ****P<0.00001.

UPEC adhesion to urinary tract cells is mediated by fimbrial adhesins with P and Type I pili conferring preferential binding to kidney and bladder epithelia, respectively^14^. A UPEC strain that spontaneously lacks both P- and type 1-pili (strain 83972) exhibited negligible binding to human or mouse tubular cells, precluding the analysis of cilia-bacteria interaction (**Fig. 1C, D**)^15^. Complementation of this strain with type 1 or P-pili restored binding to human tubular cells. While both adhesins conferred ability to bind primary cilia, preference for cilia was more pronounce for type-1 pili than for P-pili (**Fig. 1C**). In contrast, P-pili did not increase the adherence of 83972 to mouse kidney epithelial cells, while type-1 pili marginally did. The ability of complemented cells to bind cilia was weaker than the one displayed by UTI89 (**Fig. 1D**). Intriguingly, the parental 83972 strain strongly bound primary cilia of dog tubular cells, despite its lack of type I- and P-pili (**Fig. 1E**). The functionality of restored adhesins was confirmed using red blood cell agglutination and yeast binding assays (**Extended Data Fig. 1A**). Together, these data establish that UPEC binds the primary cilium of renal tubular cells across mammalian species through host-specific adhesins.

Live-cell imaging revealed that cilia-bacteria interactions were typically stable, lasting over 30 minutes, with bacteria clinging to the ciliary surface similarly to insects trapped on adhesive tape (**Fig. 1F** and **Extended Data Movie 1**). We did not observe bacteria entry into the cilium, which lacks the endocytic machinery hijacked by UPEC to become internalized. Yet, some adhering bacteria induce cilia bending. This led to fast cilia motion when they detached (**Extended Data Movie 2**).

Although these *in vitro* findings suggested UPEC preferentially bind to cilia, *in vivo* data show that UPEC initially target intercalated cells in the kidney, which lack primary cilia^16^. To investigate ciliary involvement *in vivo*, we infected seven-week-old female and male C3H/HeN mice – a sensitive strain to retrograde pyelonephritis due to spontaneous vesicoureteral reflux - with UPEC and analyzed bacterial localization relative to ciliated cells 3-, 5-, and 7-days post-infection (dpi). Pyelonephritis developed in a subset of mice (**Extended Data Fig. 1B**). In those with overt pyelonephritis, bacteria-cilia interactions were observed at 5 dpi in aquaporin 2 (AQP2^+^) collecting ducts and AQP2^-^ medullary thin limbs of Henle’s loop (**Fig. 1G** and **Sup Movies 3, 4**). At this time point, infection also triggered cilia elongation with a proportional increase in the inversin compartment (ANKS6+, **Fig. 1H)**. By day 7, infected tubules displayed widespread epithelial flattening and dedifferentiation (**Extended Data Fig. 1B)**, which was associated with cilia loss (**Extended Data Fig. 1C)**. These observations indicate that UPEC interacts with the primary cilium of tubular cells both *in vitro* and *in vivo*.

### Loss of primary cilia reduce the expression of inflammatory and fibrotic transcript in a mild model of urinary tract infection

To assess the role of primary cilia in response to *E. coli in vivo*, we used a genetic murine model allowing the inducible ablation of kidney tubule cilia upon doxycycline treatment (*Ift20*^flox/flox^.Pax8.rtTta.tetO-cre; hereafter referred to as *Ift20*^Δtub^; **Fig. 2A** and **Extended Data Fig. 2A**). By day 7 post-UPEC infection, colony-forming unit (CFU) counts showed similar bacterial loads in kidney between control and cilia-deficient mice, while bladder CFU were slightly reduced in cilia ablated male mice (**Fig. 2B and Extended Data Fig. 2B)**. Being on a C57BL/6 background without spontaneous vesicoureteral reflux, the mice showed a milder phenotype than C3H/HeN. Indeed, inspection of kidney sections showed essentially normal kidneys or isolated pyelitis and only one male mice from each genotype displayed overt pyelonephritis (**Extended Data Fig. 2C**). Intriguingly, tubule deciliation modestly but significantly reduced kidney expression of the inflammatory cytokines (*Ccl5*, *Cx3cl1* and *Cxcl10*) and fibrotic markers (*Col1a1*, *Col3a1*, *Col4a1*, *Tgfb1*, *Acta2*; **Fig. 2C-G** and **Extended Data Fig. 2D-G**). Separated analysis of males and females disclose similar trends for most transcripts with the exceptions of *Ccl5* and *Tgfb1*, which were not reduced in males (**Fig. 2G** and **Extended Data Figure 2E**). Thus, cilia ablation reduced the abundance of fibro-inflammatory transcripts in kidneys in a model of urinary tract infection by UPEC.

**Figure 2.**
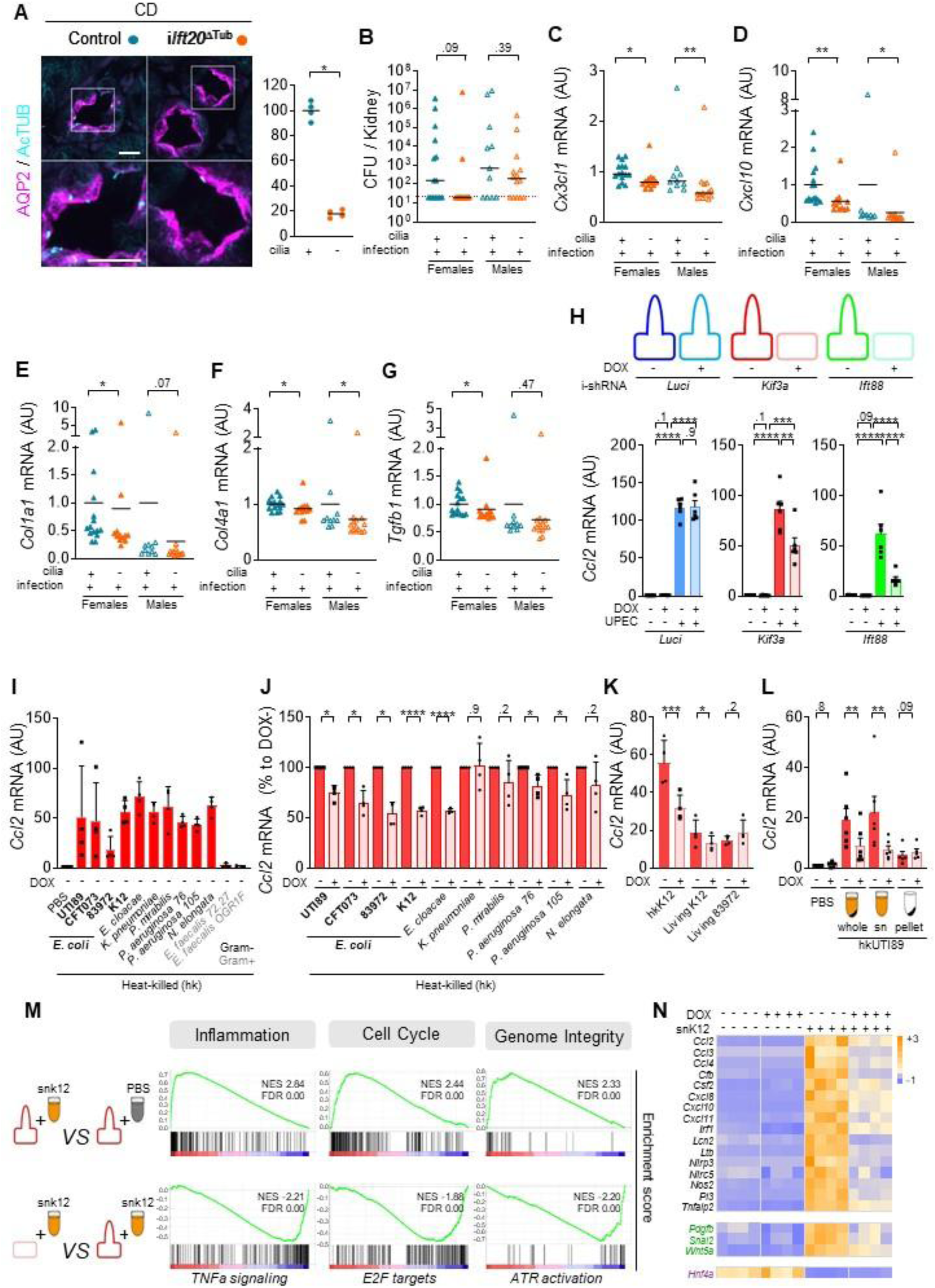
Primary cilia promote CKD-like reprogramming of tubular cells exposed to *E. coli* components. **(A)** Representative images and quantification of primary cilia (AcTUB+, cyan) in collecting ducts (CD, AQP2+, magenta) of kidneys from 8-week-old control and *Ift20*^ΔTub^ mice 2 weeks after the completion of doxycycline treatment. Scale bar: 10µm. See also **Extended Data** Fig. 2. Bars indicate mean. Each dot represents a female mouse. Mann-Whitney t-test: *P<0.05. **(B)** Colony forming units (CFU) in the kidney of 11-week-old control and *Ift20*^ΔTub^ mice, 7 days after intravesical instillation of UPEC strain UTI89. Dotted red lines depict the limit of detection of the assay, 20 CFU/bladder. **(C-G)** qPCR quantification of *Cx3cl1* (C), *Cxcl10* (D), *Col1a1* (E), *Col4a1* (F) and *Tgfb1* (G) mRNA in infected kidneys from 11-week-old control and *Ift20*^ΔTub^ mice, 7 days after intravesical instillation of UTI89. **(B-G)** Bars indicate mean. Each dot represents a kidney from a female (filled symbol) or a male (empty symbol). See also **Extended Data Fig2**. Mann-Whitney t-test: *P<0.05, **P<0.01. AU: arbitrary units. **(H)** Scheme of ciliogenesis in kidney cells (MDCK) expressing doxycycline inducible shRNA targeting *luciferase* (blue), *Kif3a* (red), or *Ift88* (green) after 10 days of culture. See also **Extended Data** Fig 3. qPCR quantification of *Ccl2* mRNA at 10 days of culture in the same cells treated for 6 hours by heat inactivated UPEC (UTI89) or vehicle (PBS). Paired one-way RM ANOVA (log-normal) with Geisser Greenhouse correction followed by Tukey test: **P<0.01, ***P<0.001, ****P<0.00001. Each dot per condition represents an independent experiment. AU: arbitrary units. **(I-J)** qPCR quantification of *Ccl2* mRNA in 10 days cultured *Kif3a* i-shRNA MDCK cells treated for 6 hours by the indicated heat killed Gram-negative (black) or -positive (grey) bacteria. Absolute induction in ciliated cells and relative effect of cilia ablation are shown in I and J, respectively. **(K)** qPCR quantification of *Ccl2* mRNA in *Kif3a* i-shRNA MDCK cells treated for 6 hours by UPEC heat-killed (hkK12) or not (living K12 and living 83972). **(L)** qPCR quantification of *Ccl2* mRNA in *Kif3a* i-shRNA MDCK cells treated for 6 hours by UPEC heat-killed (hkUTI89) in PBS (whole) and the corresponding supernatant (sn) or pellet fractions of the preparation. **(J-L)** Ratio paired *t* test: *P<0.05, **P<0.01, ***P<0.001. Each dot represents an experimental n. AU: arbitrary units. **(M)** Gene set enrichment analysis comparing selected for inflammation, cell cycle and genome integrity comparing the ciliated cells treated with *E. coli* extracts or PBS (upper pannel) or deciliated and ciliated cells treated with *E. coli* extracts (lower pannel). **(N)** Heatmap showing the Z-scores computed on the normalized counts of selected inflammatory cytokines (black), fibrogenic factors (green), and tubular differentiation (pink) markers.

### Primary cilia promote CKD-like reprogramming of tubular cells exposed to *E. coli* extracts

To gain insight into the role of the primary cilium in the response of kidney epithelial cells to UPEC, we used MDCK cell lines with inducible knockdown of genes essential for ciliary assembly and maintenance (*Kif3a* or *Ift88*) to examine whether cilia mediate epithelial responses to bacterial components (**Extended Data Fig. 3A, B**). When infected with live bacteria, MDCK cells began to detach 3–4 hours post-infection; *Ift88* knockdown, but not *Kif3a* knockdown, resulted in accelerated cell detachment and increased lactate dehydrogenase (LDH) release (**Extended Data Fig. 3C**), suggesting that IFT88 exerts protective effects against UPEC-mediated lysis through a mechanism independent of ciliogenesis. To assess whether the primary cilium participates in epithelial responses to bacterial components independently of virulence-factor-mediated cytotoxicity, we used heat-inactivated UPEC. This treatment induced a robust increase in *Ccl2* and *Cxcl10* transcripts that was significantly blunted in deciliated cells with *Kif3a* or *Ift88* depletion but not in control cells expressing a control shRNA against luciferase (**Fig. 2H** and **Extended Data Fig. 3D**). To determine if the phenotype was specifically related to cilia, we repeated the experiment three days post-seeding in doxycycline containing medium, when protein depletion is already effective but before efficient ciliogenesis (**Extended Data Fig. 3A, B, E, F**). Under these conditions, where most cells lack cilia, neither *Kif3a* nor *Ift88* silencing altered cytokine expression, indicating that the cilia themselves - not merely the associated proteins - mediate this response (**Extended Data Fig. 3G, H**).

We then tested whether other urogenital pathogens elicited similar cilia-dependent responses. We treated ciliated of deciliated (*Kif3a* depletion) MDCK with pasteurized bacteria from different strains with a urogenital tropism. Gram-positive *Enterococcus faecalis* did not increase cytokine expression (**Fig. 2I** and **Extended Data Fig. 4A**). In contrast, all tested Gram-negative *E. coli* strains strongly induced *Ccl2* and *Cxcl10* transcripts. However, cilia depletion only consistently reduced expression of both *Ccl2* and *Cxcl10* in response to *E. coli* and the closely related *E. cloacae* (**Fig. 2J** and **Extended Data Fig. 4B**). This cilia-dependent inflammatory response was not restricted to aggressive UPEC strains. Indeed, UPEC strain 83972, which does not infect human kidneys or the non-infectious K12 laboratory strain also elicited cilia-dependent inflammatory responses (**Fig. 2I, J** and **Extended Data Fig. 4A, B**).

We took advantage of these non-virulent strains to assess whether cilia mediated the response to intact living bacteria. Surprisingly, infecting tubular cells with living K12 or 83972 *E. coli* did not trigger a robust cilia-dependent response, whereas heat-inactivated preparations did (**Fig. 2K** and **Extended Data Fig. 4C**). Fractionation of heat-killed bacterial suspensions revealed that the bioactive component was not the bacteria themselves, but the supernatant in which they were heated (**Fig. 2L** and **Extended Data Fig. 4D**). These findings indicate that the primary cilium promotes the inflammatory response of kidney epithelial cells to soluble components released by injured bacteria. This may explain why cilia ablation reduced the expression of inflammatory and fibrotic transcripts in mice instillated with UPEC, in the absence of bacteria in tubular lumen (*i.e,* in the absence of pyelonephritis).

To explore the cilia-dependent transcriptional program triggered by *E. coli* extracts, we conducted bulk RNA-seq on MDCK cells with intact or ablated cilia (*via Kif3a* silencing), treated or not with heat-inactivated *E. coli* supernatant (snK12). snK12 induced a broad transcriptional response with 3724 differentially expressed genes (DEGs), of which 239 were significantly reversed by cilia ablation (**Extended Data Fig. 5A, B**). Gene Set Enrichment Analysis (GSEA) revealed that this transcriptional program encompassed three major functional modules sensitive to cilia abaltion: (i) innate immune and inflammatory signaling (*interferon-α/γ responses*, *TNFα/NF-κB*, *IL6–JAK–STAT3*, and *IL2–STAT5* signaling), (ii) cell cycle (*E2F targets*, *G2/M checkpoint*), and (iii) genome integrity (*ATR signaling*; **Fig. 2M**). Interestingly, snK12 also induced a massive downregulation of a large fraction of ciliary genes (**Extended Data Fig. 5C**), suggesting a modulation of the cilia machinery by *E. coli* components. In addition, to its effect on multiple inflammatory genes, snK12 also induced a cilia dependent increase in critical pro-fibrotic mediators such as *Pdgfb*^17,18^ or *Wnt5a*^19^ and a decrease in *Hnf4a*, a master regulator of tubular cells differentiation^20^ (**Fig. 2N**). These findings were corroborated by qPCR in both *Ift88*- and *Kif3a*-silenced cells treated with extracts derived from a distinct *E. coli* strain (uropathogenic UTI89) (**Extended Data Fig. 5D-H**). Overall, the coinciding role of cilia in the increase of fibro-inflammation, cell proliferation, DNA damage response and the reduction of tubular differentiation in response to *E. coli* extracts strikingly recapitulated the hallmarks of tubular injury that promotes CKD progression. Collectively, these results support that the primary cilium is a central regulatory node required for both immune activation and replication-associated processes.

### Cilia ablation specifically impairs the inflammatory response to ALPK1 agonists

We then sought to gain mechanistic insight into the signalling pathways regulated by the primary cilium that could be involved in response to bacterial components. Inflammatory response of cells to bacteria components usually involve the specific recognition of pathogen-associated molecular pattern (PAMPs) by innate immune receptors (pattern recognition receptors; PRR), such as toll-like receptors^21^. We analyzed how cilia ablation (*Kif3a* knock-down) impacts cell responses to typical ligands of PRR involved in the recognition of Gram^-^negative bacteria PAMPs. Surprisingly, cilia ablation did not reduce and even, in some cases, increased, cytokine expression in response to agonists of TLR1/2/4/5/6 and NOD1/2 (**Fig. 3A, B** and **Extended Data Fig. 6A**). In contrast to these negative results, cilia ablation strongly impaired cytokine induction in response to ADP-heptose (ADPh, **Fig. 3C** and **Extended Data Fig. 6B, C**). ADPh is an intermediate sugar in the biosynthesis of lipopolysaccharide, which activates inflammatory signalling by binding to ALPK1^22^, a cytosolic kinase. Once bound to ADPh, ALPK1 phosphorylates TIFA, which oligomerizes to form tifasomes. Tifasomes function as a scaffolding platform to activate inflammatory signalling through NFkB^23^. Cilia ablation also consistently reduced the induction of *Ccl2* in response to the synthetic agonist of ALPK1, DF-006^24^ (**Fig. 3D** and **Extended Data Fig. 6D**). Similarly, to the dog MDCK line, mouse derived kidney epithelial cell line mIMCD3 also responded to ADPh, *E. coli* extracts and DF-006 (**Extended Data Fig. 6E-H**).

**Figure 3.**
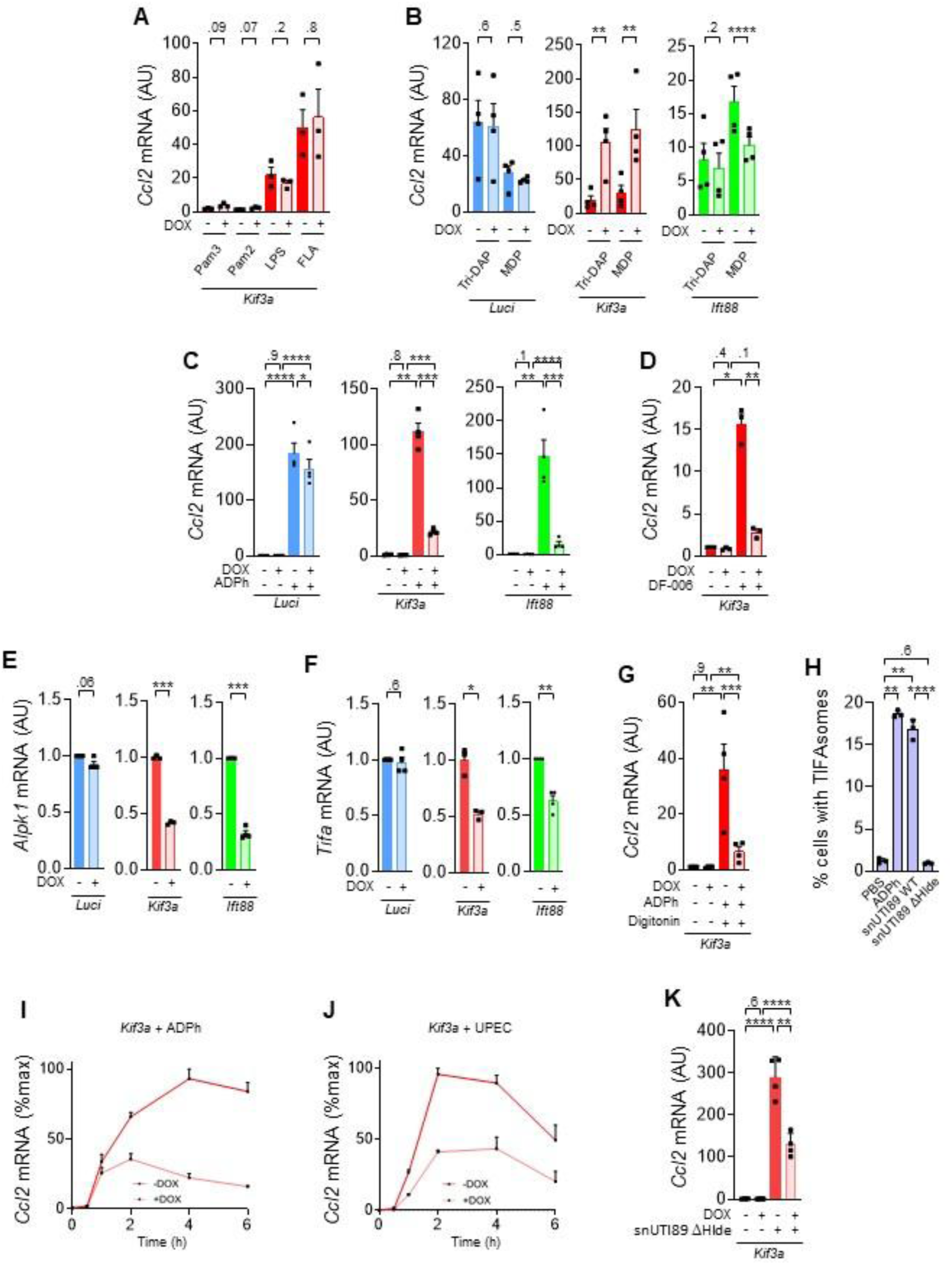
Cilia ablation specifically impairs the inflammatory response to ALPK1 agonists. **(A)** qPCR quantification of *Ccl2* mRNA in *Kif3a* i-shRNA MDCK cells treated for 6 hours with TLR-1/2 (Pam3), TRL-2/6 (Pam2), TLR-4 (Lipopolysaccharide, LPS) or TLR-5 agonist (Flagellin, FLA). **(B)** qPCR quantification of *Ccl2* mRNA in *luciferase* (blue), *Kif3a* (red) or *Ift88* (green) i-shRNA MDCK cells treated for 6 hours by NOD1 (Tri-DAP) or NOD2 agonist (MDP). **(C)** qPCR quantification of *Ccl2* mRNA in doxycycline (DOX) inducible *luciferase* (blue), *Kif3a* (red) or *Ift88* (green) shRNA MDCK cells treated for 6 hours with 13µM ADPh. **(D)** qPCR quantification of *Ccl2* mRNA in *Kif3a* i-shRNA MDCK cells treated for 4 hours with 500nM DF-006. **(E-F)** qPCR quantification of *Alpk1* (E) and *Tifa* (F) mRNA in doxycycline (DOX) inducible *luciferase* (blue), *Kif3a* (red) or *Ift88* (green) shRNA MDCK cells. **(G)** qPCR quantification of *Ccl2* mRNA in *Kif3a* i-shRNA MDCK cells treated for 30min with 13µM ADPh in presence of digitonin, and stopped 5 hours and a half later. **(H)** Percentage of cells with TIFAsome formation in HeLa cells upon ADPh, UPEC heat-killed supernatant stimulation from WT (snUTI89 WT) or lacking *Hlde* (sn UTI89 ΔHlde), an essential enzyme to generate ADPh. **(I-J)** qPCR quantification of *Ccl2* mRNA expression (expressed in percentage of the maximum value) depending on treatment duration (0min, 30min, 1h, 2h, 4h or 6h) after ADPh (I) and UPEC (J) stimulation of *Kif3a* i-shRNA MDCK cells. **(K)** qPCR quantification of *Ccl2* mRNA in *Kif3a* i-shRNA MDCK cells treated for 6 hours with UPEC heat-killed supernatant lacking *Hlde* (sn UTI89 ΔHlde). **(A-B, E-F)** Ratio paired *t* test: *P<0.05, **P<0.01, ***P<0.001, ****P<0.00001. Each dot represents an experimental n. AU: arbitrary units. **(D, G-H, K)** Paired one-way RM ANOVA (log-normal) with Geisser Greenhouse correction followed by Tukey test: *P<0.05, **P<0.01, ***P<0.001, ****P<0.00001. Each dot represents an experimental n. AU: arbitrary units.

We next investigated the mechanisms by which cilia ablation could reduce ALPK1 signalling. We found that deciliation induced a marked reduction in the abundance of *Alpk1* and *Tifa* mRNA suggesting a transcriptional regulation of ALPK1 pathway by cilia (**Fig. 3E, F)**. We ruled out a major role of cilia in the intracellular import of ADPh by showing that cell permeabilization did not interfere with the effect of cilia ablation on *Ccl2* or *Cxcl10* expression (**Fig. 3G** and **Extended Data Fig. 6I**). We then confirmed the ability of *E. coli* extracts to activate ALPK1-TIFA signalling using specific cellular assay, based on the detection of TIFAsome^25^ (**Fig. 3H**). However, while both ADPh or *E. coli* extracts triggered an increase in *Ccl2* and *Cxcl10,* the kinetics did not fully overlap. Specifically, *E. coli* extract triggered a cilia dependent early peak at 2 hours, which was not the case for ADPh (**Fig. 3I, J** and **Extended Data Fig. 6J, K**). This suggested that additional compounds present in *E. coli* extract participate to the induction of *Ccl2* and *Cxcl10*. To formerly demonstrate this, we generated an UTI89 strain lacking *Hlde* (ΔHlde), an essential enzyme to generate ADPh. Extracts from this ΔHlde UTI89 did not induce TIFAsome formation (**Fig. 3H**) but still elicited cilia dependent induction of *Ccl2* and *Cxcl10* (**Fig. 3K** and **Extended Data Fig. 6L**). Collectively these results indicate that primary cilium of kidney epithelial cells specifically control the inflammatory response to ADPh and (an) other(s) unindentified PAMP(s) released by *E. coli*.

### Primary cilia loss blunts kidney fibro-inflammatory response to urinary tract obstruction

Considering that the primary cilium participated in the induction of a transcriptional program reminiscent of CKD in response to *E. coli* components and that pathogen responsive pathways such as STING^26^ or TLR4^27^ are involved in kidney deterioration in CKD, we aimed to assess the role of the primary cilium in CKD. The primary cilium is required to maintain tubule shape in the context of tubular cell proliferation, such as kidney development and regeneration after injury^28^. In these contexts, loss of cilia leads to cystic dilation of kidney tubules, which is associated with peri-cystic inflammation and fibrosis^29^. To minimize this interfering role of cilia in tubule regeneration, we focused on the unilateral ureteral obstruction (UUO) model, in which injured tubular cells are not able to regenerate tubules but persist in an injury/failed repair state fuelling CKD progression^30^. Similarly to infection, UUO leads to an increase in cilia length with a proportional increase of the inversin compartment (ANKS6+, **Fig. 4A**) and a consistent increase in the cilia-dependent fibro-inflammatory gene signature that we identified in MDCK cells treated with UPEC (**Fig. 4B**) or in mice models recapitulating a fibrotic renal ciliopathy^6^ (nephronophtisis; NPH; **Extended Data Fig. 7A**). We applied UUO to *Ift20*^Δtub^ and control mice and analyzed them at an early time point (4 days post-obstruction; dpo), when the fibroinflammatory program is already activated but has not led to a deterioration of the kidney parenchyma. Tubule deciliation caused mild kidney enlargement and tubular dilation (**Fig. 4C** and **Extended Data Fig. 7B, C**), consistent with the established role of cilia in maintaining tubular architecture. Deciliation did not alter cytokine levels in the non-obstructed kidney, but significantly suppressed the expression of NPH-related inflammatory cytokines in the obstructed kidney (**Fig. 4D** and **Extended Data Fig. 7D-N**). This reduction correlated with decreased infiltration by mononuclear phagocytes (F4/80⁺) and neutrophils (LY6B.2⁺) in cilia-depleted obstructed kidneys (**Fig. 4E, F**). Consistent with *in vitro* data, cilia ablation also dampened fibrotic *Pdgfb* expression at 4 dpo (**Fig. 4G**), and reduced kidney fibrosis (**Fig. 4H**) and fibrosis transcripts (*Col1a1*, *Acta2*, *Tgfb1*) at 14 dpo (**Extended Data Fig. 7O-Q**) - a time point where the obstructed kidney parenchyma displays severe alterations. Deciliation further blunted the progressive loss of epithelial differentiation markers of the proximal tubules (*Hnf4a*, *Lrp2*, S*lc34a1* and *Slc26a1*) and collecting duct (*Scnn1a, Aqp2*) and preserved the tubular architecture (**Figure 4I** and **Extended Data Fig. 8**). We corroborated these results in a second model of inducible cilia ablation by inactivating *Kif3a* (**Extended Data Fig. 9-11**). Together, these findings establish a previously unrecognized role of cilia in promoting fibro-inflammatory responses to urinary flow obstruction.

**Figure 4.**
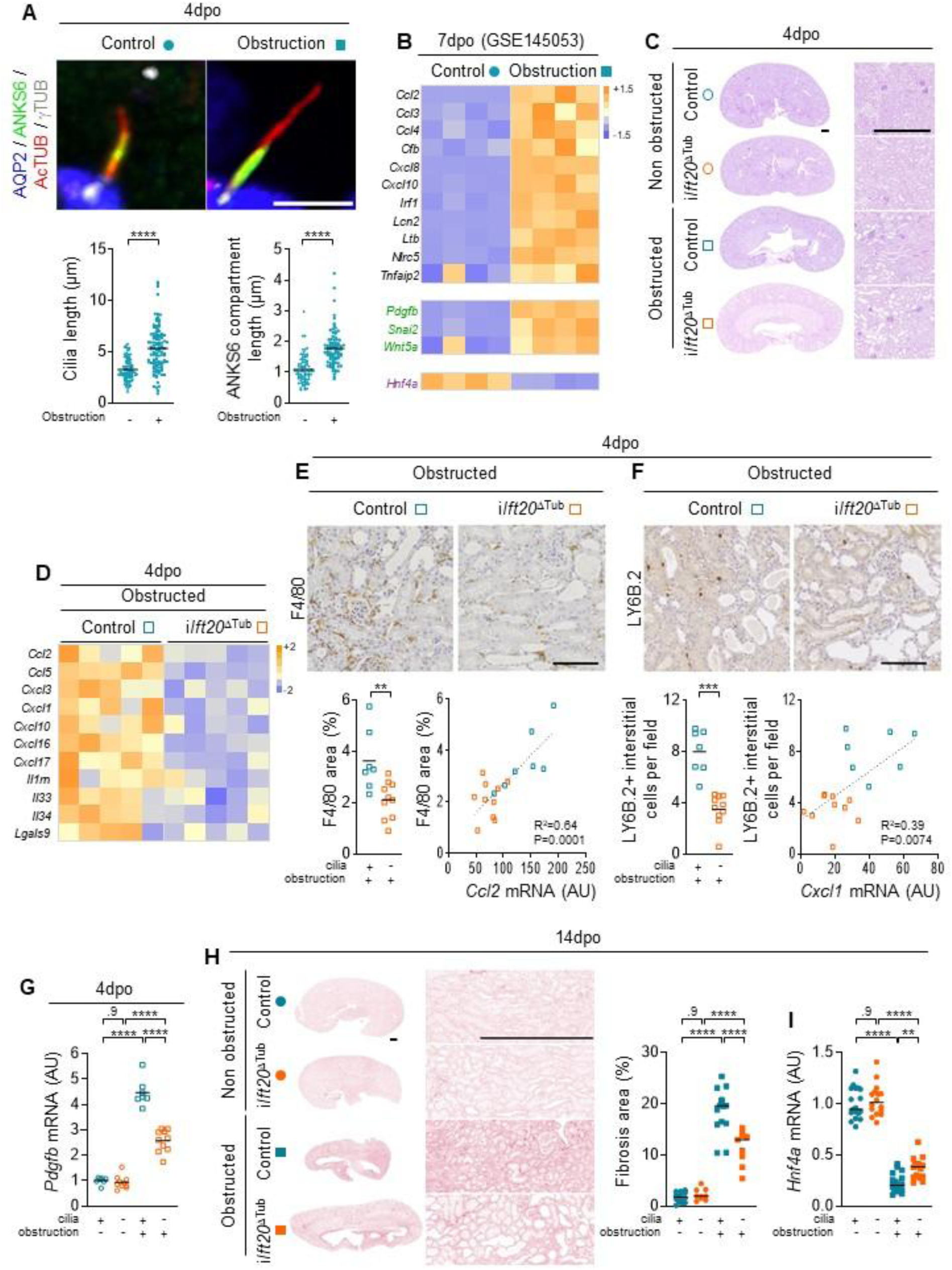
Primary cilia loss blunts kidney fibro-inflammatory response to urinary tract obstruction. **(A)** Representative images and quantification of primary cilia axoneme (AcTUB+, red; stemming from the γTUB+ basal body), and ANKS6 compartment (green) length in collecting ducts (AQP2+, blue) in control and obstructed mouse kidneys (4 days after surgery). Scale bar: 5 µm. Bars indicate mean. Each dot represents a cilium (n=3 mice per group). Mann-Whitney t-test: ****P<0.0001. **(B)** Heatmap showing the Z-scores computed on published RNAseq normalized counts (GSE145053) of the cilia-dependent fibro-inflammatory gene signature that we identified in MDCK cells stimulated with UPEC in Figure 2N**. (C)** Representative PAS staining of the obstructed and non-obstructed kidneys from 10-week-old control and *Ift20*^ΔTub^ mice, four days after surgery. Scale bar: 0.5mm. (**D**) Heatmap showing the Z-scores computed on qPCR quantifications of the indicated inflammatory transcripts in the obstructed kidneys from 10-week-old control and *Ift20*^ΔTub^ mice, four days after surgery (see also **Extended Data** Fig. 7). **(E)** Representative images and quantification of F4/80 (mononuclear phagocytes) staining and linear regression with *Ccl2* mRNA level in obstructed kidneys from 10-week-old control and *Ift20*^ΔTub^ mice, four days after surgery. Scale bars: 100 µm. **(F)** Representative images and quantification of LY6B.2 (neutrophils) staining and linear regression with *Cxcl1* mRNA level in obstructed kidneys from the same animals. Scale bars: 100 µm. **(G)** qPCR quantification of *Pdgfb* mRNA in obstructed and non-obstructed kidneys from 10-week-old control and *Ift20*^ΔTub^ mice, four days after surgery. **(H)** Representative images and quantification of picrosirius stained fibrotic area in the obstructed and non-obstructed kidney sections from 11-week-old control and *Ift20*^ΔTub^ mice, fourteen days after surgery. Scale bar: 0.5 mm. **(I)** qPCR quantification of *Hnf4a* mRNA in kidneys from 11-week-old control and *Ift20*^ΔTub^ mice, fourteen days after surgery (see also **Extended Data** Fig. 7). **(E-F)** Mann-Whitney t-test: **P<0.01, ***P<0.001. **(G-I)** One-way ANOVA followed by Tukey test: **P<0.01, ****P<0.00001. AU: arbitrary units. **(E-I)** Bars indicate mean. Each dot represents female (open symbol) or a male (close symbol) mouse.

### ALPK1 phamacologic and genetic inhibition reduces kidney fibro-inflammation in response to ureteral obstruction in rats

To directly investigate if ALPK1 activation promotes CKD progression, we took advantage of a genetic rat model allowing the invalidation of *Alpk1* and submitted these rats to UUO. Genetic inhibition of *Alpk1* reduced obstructed kidney fibrosis in rats in comparison to control rats (**Fig. 5A, B**). We further validated these results using rats submitted to UUO and treated with the ALPK1 inhibitor DF-003^31^ - an orphan drug to treat patients with ROSAH (Retinal dystrophy, Optic nerve edema, Splenomegaly, Anhidrosis, and Headache) syndrome which is caused by activating mutation in ALPK1^32^- or its vehicle. ALPK1 inhibition reduced kidney fibrosis (**Fig. 5C, D**) as well as the expression of inflammatory (*Ccl2*, *Cxcl1*, *Cxcl10*) and fibrotic (*Col1a1*, *Tgfb1*, *Acta2*) transcripts (**Fig. 5E-J**).

**Figure 5.**
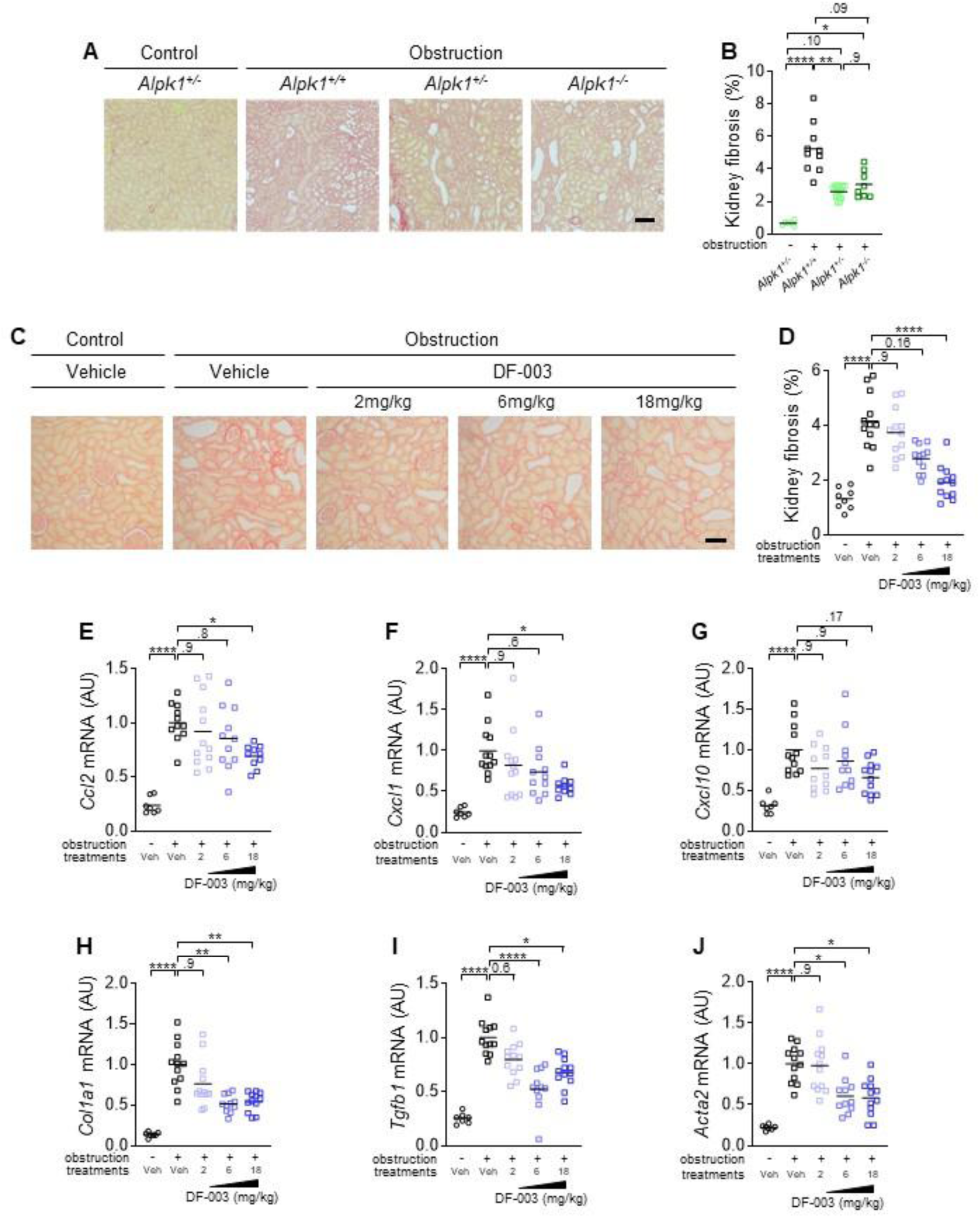
ALPK1 pharmacologic and genetic inhibition reduces kidney fibro-inflammation in response to ureteral obstruction in rats. **(A-B)** Representative images (A) and quantification (B) of picrosirius stained fibrotic area from 14-day-obstructed and non-obstructed kidneys of *Alpk1* wild-type (*Alpk1*^+/+^), heterozygous (*Alpk1*^+/-^) or knock-out (*Alpk1*^-/-^) rats**. (C-D)** Representative images (C) and quantification (D) of picrosirius stained fibrotic area from 7-day-obstructed and non-obstructed kidneys of SD rat who were dosed daily with vehicle (0.1%MC) or 1.75 (2), 6.12 (6), or 17.45 (18) mg/kg ALPK1 inhibitor DF-003, through oral gavage for 7 consecutive days. Scale bar: 150µm. **(E-J)** qPCR quantification of *Ccl2* (E), *Cxcl1* (F), *Cxcl10* (G), *Col1a1* (H), *Tgfb* (I) and *Acta2* (J) mRNA in obstructed and non-obstructed kidneys from SD rats treated with vehicle or increasing doses of DF-003, seven days after obstruction. Bars indicate mean. Each dot represents one rat. **(B, D-J)** Each dot represents a rat. One-way ANOVA with Brown-Forsythe and Welch tests followed by Dunettes *post hoc* test: *P<0.05, **P<0.01, ****P<0.0001.

### Human kidney epithelial cells display a parallel induction of ciliary and inflammatory genes in chronic injury and respond to ALPK1 agonists

We then assessed whether our results could be relevant for human. As a starting point, we correlated the expression of ciliary genes with tubular cell state at a single cell level, leveraging available human datasets^33^. We observed that expression of ciliary genes was up-regulated in injured tubular cells in comparison to non-injured cells (**Fig. 6A, B**). Strikingly, ciliary gene expression correlated with NF-κB target gene expression, which was most pronounced in injured tubular cells (**Fig. 6C** and **Extended Data Fig. 12A**). Thought we failed to detect robust expression of ALPK1 and TIFA in injured tubular cells, this was also the case for other important PRR involved in CKD progression (**Extended Data Fig. 12B)**.

**Figure 6.**
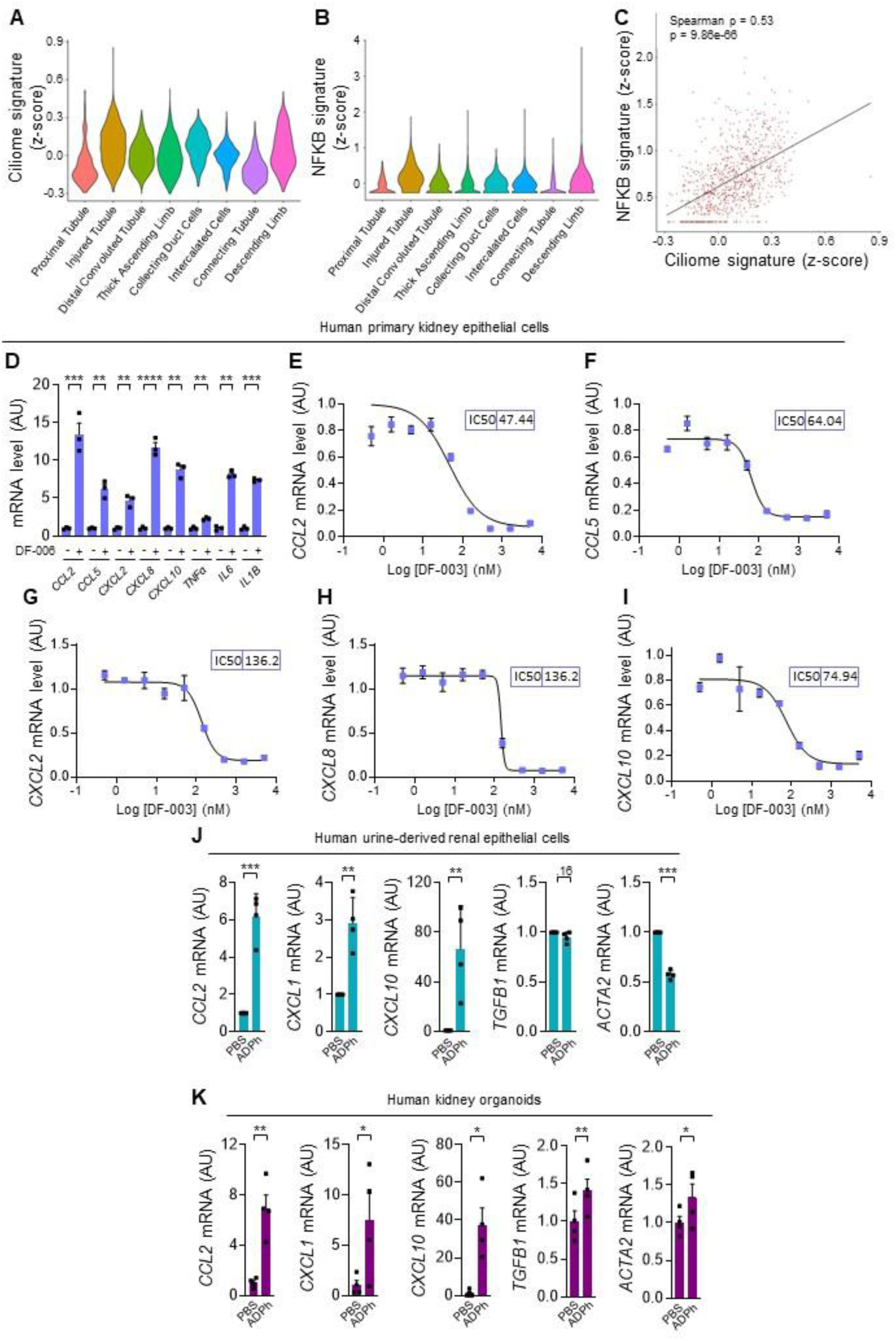
Human kidney epithelial cells display a parallel induction of ciliary and inflammatory genes in chronic injury and respond to ALPK1 agonists. **(A-C)** Single-cell RNA sequencing data from the Kramann human kidney atlas were analyzed. Signature scores were computed as the mean z-score of log-normalized expression values across retained signature genes (ciliary: Syscilia ciliome database; NF-κB: BIOCARTA_NFKB_PATHWAY gene set). (A) Violin plots showing the distribution of ciliome signature scores across eight tubular cell types. (B) Violin plots showing the distribution of NF-κB pathway signature scores across the same tubular populations. (C) Scatter plot of ciliome versus NF-κB signature scores in injured tubular cells, with Spearman correlation coefficient and linear regression fit (gray line) see also **Extended figure data 12. (D-I)** qPCR quantification of *CCL2, CCL5, CXCL2, CXCL8* and *CXCL10* mRNA in human primary kidney epithelial cells treated for 4 h with DF-006 (100 nM) (D), or pre-treated for 2 h with increasing doses of DF-003 (E-I). See also **Extended figure data 12**. **(J)** qPCR quantification of *CCL2*, *CXCL1, CXCL10, TGFB1* and *ACTA2* mRNA in human urine-derived renal epithelial cells treated or not for 6 hours with ADPh (13µM). **(K)** qPCR quantification of *CCL2*, *CXCL1*, *CXCL10*, *TGFB1* and *ACTA2* mRNA in human kidney organoids treated or not with 13µM ADPh during 3 days. **(D, J-K)** Each dot represents an experimental n. AU: arbitrary units. Ratio paired *t* test: *P<0.05, **P<0.01, ***P<0.001, ****P<0.00001.

Therefore, we assessed whether human tubular cells have a functional ALPK1 pathway. We found that ALPK1 agonist DF-006 induced inflammatory cytokine expression in human primary kidney epithelial cells, that could be inhibited by the ALPK1 inhibitor DF-003 (**Fig. 6D-I** and **Extended Data Fig. 12C-E**). Similarly, ADPh treatment of human urine-derived renal epithelial cells (URECs) and iPS-derived kidney organoids prompted a fibro-inflammatory response (**Fig. 6J-K)**.

## DISCUSSION

Our findings reveal an unanticipated role for the primary cilium of kidney epithelial cells as a UPEC-binding structure that calibrates the epithelial innate-immune response to specific *E. coli* components, with direct relevance to CKD.

We first show that primary cilia of tubular cells from multiple mammalian species avidly bind uropathogenic E. coli (UPEC), and that this binding is mediated by species-specific adhesins — with Type 1 and, to a lesser extent, P fimbriae supporting attachment to the cilia of human kidney tubular cells. Binding is not followed by bacterial endocytosis, at least over the timescales examined — consistent with the absence of endocytic machinery in the ciliary membrane. The functional consequence of this capture remains to be formally established, but we propose that, at low bacterial loads, the cilium acts as a molecular “flypaper”: it tethers bacteria to a surface they cannot exploit for invasion while holding them within range of receptors that detect pathogen-associated molecular patterns (PAMPs) released upon bacterial injury. In this view, ciliated principal cells and non-ciliated intercalated cells divide labour within the distal nephron — the latter providing direct antimicrobial activity through urine acidification and defensin secretion^34^, the former operating as a sentinel that converts ligand detection into a paracrine cytokine signal to recruit immune and mesenchymal effectors. Consistent with this model, cilia amplify the fibro-inflammatory transcriptional response of tubular cells to defined E. coli PAMPs.

This cilia-dependent transcriptional programme mirrors core features of tubular reprogramming in CKD, including the induction of pro-inflammatory and profibrotic cytokines and the repression of HNF4α, a master regulator of tubular transport. These in vitro observations were recapitulated in *vivo*, where cilia ablation attenuated the fibro-inflammatory response to both urinary tract infection and ureteral obstruction. In obstructive models, cilia loss reshaped the kidney response in a reproducible and bidirectional manner: deciliation exacerbated tubular dilation but consistently blunted inflammation, dedifferentiation and fibrosis. Using independent deciliation strategies, we confirmed that these effects reflect bona fide ciliary functions rather than cilia-independent activities of intraflagellar transport components such as IFT88, which can modulate immune signalling in non-ciliated cells^35^.

Tubular dilation accompanied by inflammation is a well-documented consequence of cilia loss during kidney development^28,29^. Our data identify a contrasting role: in the adult kidney cilia can actively promote fibro-inflammation. Across disparate triggers — E. coli extracts, ADP-heptose, and ureteral obstruction — the fibro-inflammatory output is amplified by an intact cilium, echoing the cilia-dependent pathogenic signalling described in autosomal dominant polycystic kidney disease^5,36^.

This coupling of antibacterial defence to tissue remodelling may reflect an evolutionarily conserved adaptation. Ureteral obstruction is life-threatening when complicated by infection, and selection may have favoured cilium-mediated responses that downregulate energetically costly transport programmes while recruiting immune sentinels; the ensuing vascular rarefaction and matrix deposition may act, in the short term, as a secondary barrier limiting bacterial dissemination from the tubular lumen to the bloodstream. The same programme, chronically engaged, would plausibly drive the scarring that underlies CKD.

Mechanistically, our data implicate ALPK1 as an important relay of cilia-dependent inflammatory signalling. Cilia loss markedly attenuated ALPK1 pathway activation, and ALPK1 ligands induced fibro-inflammatory programmes in human tubular cells and iPSC-derived kidney organoids. In parallel, ciliary gene expression rises in human tubular cells that acquire a fibro-inflammatory phenotype in CKD. Recent work points to the gut microbiota as a systemic source of ADP-heptose, particularly with aging, where dysbiosis and increased intestinal permeability elevate circulating levels of this metabolite and promote clonal haematopoiesis— itself associated with CKD. Whether ADP-heptose directly sustains tubular fibro-inflammation in the aged kidney remains to be tested. ALPK1 is unlikely to be the sole cilium-coupled trigger of this programme: an E. coli ΔhldE strain, unable to synthesize ADP-heptose, still elicited a cilia-dependent inflammatory response, implicating additional PAMPs whose signalling is amplified by the cilium independently of ALPK1.

Several questions remain open. Although inducible, tubule-specific deciliation established that cilia shape the kidney’s fibro-inflammatory response, we did not fully resolve their contribution to antibacterial defence itself: UPEC infection is mild in C57BL/6 mice, and more permissive models of ascending infection will be needed to determine whether ciliary capture of bacteria alters the kinetics of bacterial clearance or dissemination. A second open question concerns the nature of the ciliary contribution to signalling — whether cilia directly sense bacterial PAMPs at the axonemal membrane or instead amplify signals first perceived elsewhere in the cell. In the case of ALPK1, the cilium appears to act, at least in part, by modulating ALPK1 and TIFA expression. Yet the ALPK1-independent component uncovered by our ADP-heptose-deficient strain points to at least one additional PAMP, amplified through the cilium, whose identity remains to be defined. Finally, our human data are correlative: the coupled ciliary–inflammatory transcriptional signature observed in CKD tubular cells is consistent with the mechanism we propose but does not, on its own, establish causality in patients.

Together, these findings identify the primary cilium as a previously unappreciated node coupling epithelial surveillance of luminal bacteria to downstream tissue-remodelling programmes. This dual function positions cilia fibro-inflammatory signalling as a candidate therapeutic target in CKD.

## MATERIAL AND METHODS

### Mice

Mice were maintained under specific pathogen-free conditions at a constant ambient temperature on a 12-hour light/dark cycle, with *ad libitum* access to food and water in an enriched environment. Breeding and genotyping were performed following standard procedures. All mice were on a C57BL/6 (mixed J/N) background.

*Kif3a*^flox/flox^ mice (Kif3a^tm1Gsn^, C57BL/6 background, kindly provided by Peter Igarashi, Stony Brook University) and *Ift20*^flox/flox^ mice (Ift20^tm1.1Gjp^, kindly provided by Gregory Pazour, UMass Chan Medical School) were crossed with Pax.8rtTA (Tg[Pax8-rtTA2S*M2]1Koes, Jackson Laboratory) and TetO.Cre (Tg[tetO-cre]1Jaw, Jackson Laboratory) mice to generate inducible, tubule-specific *Kif3a* knockout (*Kif3a*^ΔTub^) and *Ift20* knockout (*Ift20*^ΔTub^) lines.

Doxycycline hyclate (Abcam, ab141091; 2 mg/mL) supplemented with 5% sucrose and protected from light was provided in drinking water from postnatal day 28 to 42 to induce Cre-mediated recombination. Littermates lacking either TetO.Cre or Pax8.rtTA served as controls. Both sexes were included in all experiments, and all studied animals were included in the analyses.

For bacterial infection experiments, 10-week-old control or *Ift20*^ΔTub^ mice were anesthetized by intraperitoneal injection of ketamine (100 mg/kg) and xylazine (5 mg/kg) and catheterized transurethrally, and infected with 10^8^ CFU of UTI89-GFP-amp^R37^ in 50 μl of phosphate-buffered saline (PBS) as previously described^34^. To ensure propagation of bacteria into the kidney, two rapid instillations are performed 3 hours apart. Animals were sacrificed by cervical dislocation after isoflurane inhalation, and bladders and kidneys were aseptically collected. The bladder and right kidney were homogenized in 1 ml of cold sterile PBS, serially diluted, and plated on LB agar plates with carbenicillin (100µg/ml) to determine CFU. The left kidney was longitudinally cut in 2 equal pieces. One part was fixed in 4% PFA and the other part was flash-frozen and stored in liquid nitrogen.

For unilateral ureteral obstruction (UUO) experiments, 8-week-old control, *Kif3a*^ΔTub,^ or *Ift20*^ΔTub^ mice were used as previously described^38^. Briefly, mice were anesthetized by intraperitoneal injection of ketamine (160 mg·kg⁻¹) and xylazine (8 mg·kg⁻¹) and received buprenorphine (0.1 mg·kg⁻¹) for postoperative analgesia. Under a dissecting microscope, a left flank incision was made to expose the ureter, which was ligated with nonabsorbable 6/0 black braided silk suture. Mice were euthanized by intraperitoneal injection of ketamine 160 mg·kg⁻¹ and xylazine 8 mg·kg⁻¹, followed by cervical dislocation upon confirmation of deep anesthesia.

### Rats

Rat studies were approved by the Institutional Animal Care and Use Committee (IACUC) at Shanghai Yao Yuan Biotechnology Ltd. (Drug Farm) and Inotiv Westminster. Wildtype Sprague Dawley (SD) rats were acquired from Charles River Laboratories and were housed in a specific pathogen-free facility in ventilated cages on a 12 h light/dark cycle with corncob bedding, and *ad libitum* access to water and rodent chow (Keaoxieli, Beijing, China or Envigo, #8940). Rats were allowed to acclimate for at least 7 days prior to use.

To generate *Alpk1*^−/−^ rats, guide RNAs (gRNAs) (5’-TTGCTTGCACTCGTGCAGTA-3’ and 5’-AGAGACGTCACTACAAACTT-3’) targeting exon 2 and 14 of *Alpk1* transcript (XM_227715.9) were designed with an online gRNA designing tool (http://crispr.mit.edu/). *In vitro* transcribed gRNA and Cas9 mRNA were co-microinjected into rats-derived fertilized eggs. The tail-end genomic DNA of each offspring was amplified with the forward primer 5′-GGTCCTGTCATGTATTTTCATGC-3′ and the reverse primer 5′-CCATTGTGGGTTGGTTGGAGAC-3′ and Sanger sequencing was performed. One founder with ∼55kb genomic DNA deletion between exon 2 and exon 14, was backcrossed to wild-type SD rats to obtain *Alpk1*^+/−^ rats and these heterozygote rats were intercrossed to generate *Alpk1*^−/−^ homozygotes. *Alpk1*^+/+^ littermates in the offspring were used as controls.

In unilateral ureteral obstruction (UUO) study performed on *Alpk1*^+/+^, *Alpk1*^+/-^ and *Alpk1*^−/−^ rats, six-week-old male rats weighing 150-200 g were used. On day 0, 6 *Alpk1*^+/-^ rats underwent sham (no occlusion) surgery; remaining *Alpk1*^+/-^ rats and all *Alpk1*^+/+^, and *Alpk1*^−/−^ rats underwent permanent right UUO surgery.

To evaluate DF-003^31^ efficacy in UUO model, ∼8-week-old SD male rats, weighing ∼170-200 g, were placed into weight-matched treatment groups. Group 1 underwent sham (no occlusion) surgery and received Vehicle; Group 2 underwent permanent right UUO and received Vehicle; Groups 3-5 underwent permanent right UUO and were received DF-003 once a day (QD) at doses of 2, 6 and 18 mg/kg body weight, respectively. Surgical manipulations were performed using heat sterilized instruments and aseptic surgical technique. Immediately following completion of surgery, all rats received a single subcutaneous (s.c.) injection of 0.12 mg/kg buprenorphine and recovered in clean, heated cages before returning to normal vivarium conditions. All dosing was given through oral gavage from day 0 through day 7 in volumes equal to 5 mL/kg body weight. On day 7, all rats were anesthetized with isoflurane. Each right kidney was collected, placed in ice-cold 0.9% NaCl, decapsulated, blotted off excess saline, and weighed. One mid-transverse short-axis section (∼ 3mm) from each right kidney (sham or obstructed) was immersion-fixed in 10% neutral buffered formalin (NBF). All remaining renal cortical tissue from each right kidney was collected, flash-frozen in liquid nitrogen, and stored at -80°C until cryogenically powdered.

### Histopathology and Morphometric Analysis

Mouse kidneys were fixed in 4% paraformaldehyde for 24 hours at 4°C and embedded in paraffin. Four-µm sections were stained with Periodic Acid–Schiff (PAS) or picrosirius red.

Whole-slide images were acquired using a NanoZoomer 2.0 scanner (Hamamatsu) equipped with a 20×/0.75 NA objective and analyzed with NDPview software (Hamamatsu).

For fibrosis quantification, picroSirius red stained area was measured with FIJI software^39^ from full sized mouse kidney images and visualized as the percentage of stained surface to total kidney section area.

Rat fixed kidneys were paraffin embedded and had 1 section (5 µm) obtained at each of 3 anatomically-distinct depths. Sections of 5 µm were stained with picrosirius red. Images (n = 10/section) of picrosirius red-stained renal tissue were obtained in a blinded manner using a Zeiss AxioImager.A2 microscope at 200x magnification (enough to obtain a representation of ∼60 – 70% of the renal cortical area). Quantitative histological image analysis of picrosirius red -stained tissue sections was performed using color spectral segmentation methods. Fibrosis percentage was expressed as the average positive stain across sampled images.

### Immunohistochemistry (IHC) and Immunofluorescence (IF)

Four-µm (all staining except cilium) or 7µm (cilium staining only) sections of paraffin-embedded mouse kidneys were used for IHC and IF. Cultured cells grown on glass coverslips or Ibidi® slides were used only for IF. Mouse kidney sections were first submitted to paraffin removal and antigen retrieval treatment (10 mM Tris pH9 – 1 mM EDTA – 0.05% Tween-20 or Dako REAL Target Retrieval Solution #S2031). For IHC, we performed 3% H2O2 (Sigma, #95321) and avidin/biotin blocking (Vector, #SP-2001) steps. Sections were then blocked with TBST – 10% FBS for 20 minutes and incubated overnight at 4 °C with the indicated primary antibodies. The next day, sections were incubated with appropriate secondary antibodies and a final step of 3,30-diaminobenzidine-tetrahydrochloride (DAB) revelation was performed for IHC. Antibodies used are listed in Supplementary Table 1. For IHC, full size images were recorded using a whole slide scanner NanoZoomer (Hamamatsu) equipped with a 20×/0.75 NA objective and coupled to NDPview software (Hamamatsu). For IF, confocal images were acquired using a Spinning Disk microscope (40X NA1.3 or 63X NA1.4 oil immersion, Zeiss). All quantifications were performed using FIJI software. For macrophage quantification, we measured the percentage of stained F4/80 area in all injured areas of the kidney. For neutrophil (Ly6B staining) quantification, we counted manually the number of foci in the whole kidney section. Foci were defined as 2 or more neutrophils surrounding a tubule. For these 2 quantifications, glomerular and intra-tubular positive stainings were removed from the analysis. For IF on cultured cells, slides were subjected to antigen retrieval, followed by avidin/biotin blocking or permeabilization. Samples were incubated with primary antibodies, followed by Alexa Fluor–conjugated secondary antibodies, fluorescently labelled lectins, and Hoechst 33342 (Thermo Fisher).

### Quantitative Real-Time PCR (qRT-PCR)

Total RNAs were obtained from mouse kidneys or cultured cells using RNeasy Mini Kit (Qiagen) or the NucleoSpin® RNA Kit (Macherey-Nagel), and reverse transcribed using High-Capacity cDNA Reverse Transcription Kit (Thermo Fisher Scientific) according to the manufacturer’s protocol. Quantitative PCR were performed with iTaq™ Universal SYBR® Green Supermix (Bio-Rad) on a CFX384 C1000 Touch (Bio-Rad). Each biological replicate was measured in technical duplicates. *Rpl13*, *Hprt, Gapdh, Sdha, Tbp* and *Ppia* were used as normalization controls^40^. The primers used for qRT-PCR are listed in Supplementary Table 2. Heatmaps displaying Z-scores computed on the expression levels of the identified cytokines, measured by qPCR, were generated using excel.

To extract total RNA from human primary kidney epithelial cells (Cell Biologics #H6621), supernatants were removed and 500 μL RNAiso Plus (Takara #9109) was added to each well.

RNA was extracted from the cells following the reagent manufacturer’s instructions and finally dissolved in RNase-free water.

To extract total RNA from rat kidneys, cryo-genically powdered tissues were lysed with TRI reagent® (Sigma-Aldrich T9424) before mixing with chloroform. RNA in the aqua phase was precipitated with isopropanol and ethanol, and finally dissolved in RNase-free water. Blood RNA was extracted by QIAamp RNA Blood Mini Kit (Qiagen 52304). RNA from human primary kidney cell and rat kidney was reverse-transcribed using HiScript Q RT SuperMix (Vazyme Biotech R122-01, Nanjing, China) with random primers/Oligo(dT) primer mix. The synthesized cDNA was used for qRT-PCR analysis with AceQ qPCR SYBR green master mix (Vazyme Q111-02) and gene-specific primers (Supplementary Table 2) on a QuantStudio™ 7 Flex Real-Time PCR System (Thermo Fisher). Gene expression values were calculated by comparative ΔΔCt method after normalization to *Ubc* (rat) and *GAPDH* (human) internal control and presented as fold changes over the mean value of UUO-Vehicle group (rat kidney) or Vehicle (human kidney cell).

### Bacterial Strains and Culture Conditions

All bacterial strains (Supplementary Table 3) were stored in Luria–Bertani (LB) or Brain Heart Infusion (BHI; for *Neisseria elongata*) broth with 10% glycerol at –80°C. Cultures were streaked on LB agar plates and, unless otherwise indicated, single colonies were inoculated into 5–10 mL of LB (or BHI for *N. elongata*) and grown overnight at 37°C with shaking. Specifically for mouse infections, bacteria were grown standing in 10 mL LB containing appropriate antibiotics. For larger volumes, 5 mL of overnight culture was transferred after 2 hours into 100–200 mL of LB and grown overnight at 37°C. When required, antibiotics were used at the following concentrations: kanamycin (50 µg/mL), chloramphenicol (34 µg/mL), and carbenicillin (100 µg/mL).

### Bacterial heat-inactivation and Extract Preparation

For bacterial pasteurization, 10⁹ (heat inactivated bacteria) or 10¹⁰ (for concentrated extracts) bacteria were resuspended in PBS and incubated at 60°C for 1 hour in a water bath. Samples were centrifuged at 5,000 × g for 5 minutes, and the supernatant was collected, filtered through a 0.2 µm filter (Corning), and stored at –80°C.

### Cell Culture

Madin–Darby canine kidney (MDCK; kind gift of Prof. Kai Simons, MPI-CBG, Dresden, Germany) were cultured in Dulbecco’s Modified Eagle Medium (DMEM) supplemented with 10% fetal bovine serum (FBS) and 1% penicillin–streptomycin (Sigma). To generate MDCK cell lines for tetracycline-inducible knockdown of target genes, a lentivirus-based transduction system (pLVTH) was used as previously described^5^. Expression of the shRNA was induced by tetracycline (Abcam, ab141091) at 1µg/mL 3 hours after seeding and changed every 2 days for 10 days. Venus-STRADb expressing MDCK lines was generated with a retroviral transduction system (pLXSN) as previously described^5^.

Mouse inner medullary collecting duct (mIMCD3) cells were maintained in a 1:1 mixture of DMEM high-glucose pyruvate (Gibco, 41966052) and F-12 Nutrient Mixture (Gibco, 21765037) supplemented with 10% FBS and 1% penicillin–streptomycin.

HK2 cells were cultured in a 1:1 mixture of DMEM high-glucose pyruvate and F-12 Nutrient Mixture supplemented with 1% FBS, 1% penicillin–streptomycin, 5 μg/mL insulin, 5 μg/mL transferrin, 50 nmol/L selenium (ITS; Thermo Fisher), 10 mM HEPES, 5 pM triiodo-L-thyronine (Sigma, T6397), 10 ng/mL epidermal growth factor (Sigma, E9644), 3.5 µg/mL ascorbic acid (Sigma, A4544), 25 ng/mL prostaglandin E1 (Sigma, P5515), and hydrocortisone (Sigma, H0396).

hTERT-RPE1 cells were cultured in DMEM supplemented with 10% FBS and 1% penicillin–streptomycin.

HeLa cells (ATCC) were cultured in DMEM supplemented with 10% FBS, 2 mM GlutaMAX-1, 100 μg/mL streptomycin and 100 U/mL penicillin at 37 °C under 5% CO2. HeLa cells stably expressing GFP-TIFA were previously described^41^.

Human primary kidney epithelial cells (Cell Biologics #H6621) were cultured in complete human endothelial cell medium (H6621) with a supplement kit (H6621- Kit). Cells were plated at a density of 80,000 cells/well in a 24-well plate precoated with Collagen Type IV (Sigma #C5533). The cells were maintained at 37°C in a humidified incubator with 95% atmospheric air and 5% CO2 overnight. The cell culture medium was refreshed the next day.

### L-Lactate dehydrogenase (LDH) quantification

LDH release was quantified using LDH-Glo Cytotoxic Assay (Promega) following the manufacturer’s instructions.

### TIFAsomes quantification

HeLa cell expressing GFP-TIFA were fixed in 4% paraformaldehyde and incubated with Hoechst (Life Technologies, H3570) diluted in 0.2% saponin for 45 min. Images were acquired with an ImageXpress Micro (Molecular Devices, Sunnyvale, USA). Quantification of cell fraction with TIFAsomes was performed using MetaXpress as previously described^41^. Each data point corresponds to triplicate wells, and more than nine images were taken per well.

### Agglutination test on yeast and erythrocytes

To detect PapG- and FimH-mediated agglutination, bacteria were grown overnight in static LB cultures at 37°C to induce pili expression. Agglutination was assessed using human red blood cells (8% vol/vol in PBS) or a baker’s yeast suspension (5% vol/vol in PBS). Human erythrocytes were obtained from remnant blood samples originally collected for routine plasma parathyroid hormone (PTH) measurement in the Department of Physiology (Hôpital Necker – Enfants Malades, AP-HP, Paris). The use of remnant biological samples was conducted in accordance with French regulations governing the secondary use of samples collected for clinical care purposes (Article L.1211-2 of the French Public Health Code). Non-opposition to the potential reuse of residual samples for research was obtained from all patients at the time of sampling.

### Scanning electron microscopy

MDCK samples were fixed in 2.5% glutaraldehyde in 1X Hepes buffer O/N at 4°C, postfixed for 1h in 1% osmium. Then, samples were dehydrated through a graded ethanol series followed by critical point drying with CO2 (CPD300, Leica). Dried specimens were gold/palladium sputter-coated with a metal coater (ACE600, Leica). The samples are imaged in a JEOL IT700HR field emission scanning electron microscope.

### Western Blotting

MDCK cells were lysed in RIPA buffer (20 mM Tris pH 8.0, 160 mM NaCl, 1 mM EDTA, 1 mM EGTA, 1 mM DTT, 0.1% SDS, 1% NP-40, 1% sodium deoxycholate, 1% Triton X-100) using a 20G needle. Lysis buffers were supplemented with 1 mM Na₃VO₄, 50 mM NaF, 5 mM β-glycerophosphate, and a protease inhibitor cocktail (Roche). Protein concentration was determined using the Pierce BCA Protein Assay Kit (Thermo Fisher Scientific). Equal protein amounts were resolved on 4–15% Mini-PROTEAN TGX™ gels (Bio-Rad) under reducing conditions, transferred to membranes, and probed with specific primary and secondary antibodies. Bands were visualized on film, and densitometry was performed using LabImage 1D L340 software.

### Cell Infections

MDCK (200,000 cells/cm²), mIMCD3 (50,000 cells/cm²), HK2 (50,000 cells/cm²), and hTERT-RPE1 (60,000 cells/cm²) cells were seeded on glass coverslips (MDCK, mIMCD3, hTERT-RPE1) or Ibidi® 8-well slides (HK2). After 9 days (MDCK), 3 days (mIMCD3, HK2), or 2 days (hTERT-RPE1) of culture, the medium was replaced with antibiotic- and serum-free medium for 24 hours before infection with 2×10⁶ bacteria for 2 hours. Cells were then washed with PBS and fixed in 4% paraformaldehyde for 10 minutes.

For live-cell imaging, MDCK cells expressing Venus– STRADb (a cilium-localized fusion protein)^5^ were seeded (200,000 cells/cm²) on Ibidi® 8-well slides and cultured for 9 days. After 24 hours in antibiotic- and serum-free DMEM, cells were infected with 2×10⁶ UPEC UTI89-RFP bacteria for 2 hours in EM buffer (20 mM NaCl, 7 mM KCl, 1.8 mM CaCl₂, 0.8 mM MgCl₂, 5 mM glucose, 25 mM HEPES, pH 7.3). Cells were washed in EM buffer before time lapse microscopy. Imaging was carried out on a Nikon Ti-E inverted microscope equipped with a Perfect Focus System (PFS), a spinning disk confocal system (CSU-W1, Yokogawa), and an ORCA flash 4.0 camera (Hamamatsu). Time-lapse imaging was performed using a CFI S Plan Fluor ELWD 40x/0.60 air immersion objective (MRH08430, Nikon) or with a CFI Plan Apo VC 60X/1.2 water immersion objective (MRD07602, Nikon) or CFI Plan Apo 60X/1.4 oil immersion objective (MRD01605, Nikon). Time lapses were taken every three minutes, and automated acquisition of multiple positions was achieved with the NIS-Elements software (Nikon).

### Reagents

ADPh (tlrl-adph-l, InvivoGen) was dissolved in endotoxin-free water at 1mg/mL and added to the cell culture medium at 10µg/mL final concentration during 6 hours. CLI-095 (tlrl-cli95-4, InvivoGen) was dissolved in DMSO at 2.5 mM and added to the cell culture medium at 5µM final concentration during 1 hour of pre-treatment before stimulation. DF-003^31^ was 3.16-fold serially diluted and tested at 10 concentrations in triplicate. 0.08% DMSO served as vehicle control. Two hours later, 100 nM DF-006^24^ dissolved in PBS was added to the cells to activate ALPK1 and stimulate pro-inflammatory gene expression during 4 hours of DF-006 incubation. FLA (tlrl-epstfla, InvivoGen) was dissolved in sterile water at 100 µg/mL and added to the cell culture medium at 100ng/mL final concentration during 6 hours. LPS (tlrl-b5lps, Invivogen) was dissolved in endotoxin-free water at 5 mg/ml and added to the cell culture medium at 5µg/mL final concentration during 6 hours. MDP (tlrl-mdp, InvivoGen) was dissolved in sterile water at 10 mg/mL and added to the cell culture medium at 10µg/mL final concentration during 6 hours. Pam2 (tlrl-pm2s-1, InvivoGen) and Pam3 (tlrl-pms, InvivoGen) were dissolved in endotoxin-free water at 5 mg/ml and added to the cell culture medium at 5µg/mL final concentration during 1 hour of pre-treatment before stimulation. Tri-DAP (tlrl-tdap, InvivoGen) was dissolved in sterile water at 10 mg/mL and added to the cell culture medium at 10µg/mL final concentration during 6 hours. See Supplementary Table 4.

### Cilia / Bacteria Interaction Analysis

A maximum intensity Z projection has been applied to the dataset using FIJI software. Next, cilia and bacteria segmentation were performed using the ilastik v1.3.3post3^42^ workflow based on supervised shallow learning pixel classification and hysteresis thresholding of the probability map of positively labeled pixels with at least a size filter. Two general models have been performed for each dedicated staining: acTUB for cilia and RFP for bacteria. The generated output is a cilium and a bacteria label map. Finally, in a FIJI macro, we import the raw data, the cilia and bacteria labelling maps from Ilastik. We then count the total number of bacteria and the number of bacteria located at a specified distance of 0.52 µm around the cilia. To determine whether the placement of bacteria near the cilia is not a random factor, we applied horizontal and vertical symmetry to the bacterial labelling map and performed the same count on both artificial datasets. If the difference between the mean counting of the 2 flipped images and the raw image are significant, we validate that the bacteria proximity is not randomized.

For this study, X cell of Y filed on Z experiments have been analyzed.

### Bulk RNA-seq Analysis

Total RNAs were extracted and purified from MDCK using NucleoSpin® RNA Kit (Macherey-Nagel), RNA quality was verified by capillary electrophoresis using Fragment AnalyzerTM. Bulk mRNA-Seq libraries were prepared by the Imagine Genomic Core Facility (Paris, France) starting from 1µg of total RNA, using the NEBNext® Low Input RNA Library Prep Kit according to manufacturer’s guidelines. This kit generates mRNA-Seq libraries from the PolyA+fraction of the total RNA by template switch reverse transcription. The cDNA are amplified twice and the final mRNAseq libraries are not ‘oriented’ or ‘stranded’ (meaning that the information about the transcribed DNA strand is not preserved during the library preparation). The equimolar pool of libraries, assessed by Q-PCR KAPA Library Quantification kit (Roche) and with a run test using the iSeq100 (Illumina) was sequenced on a NovaSeq6000 (Illumina, S2 FlowCells, paired-end 100+100 bases, ∼50 millions of reads/clusters produced per library). After the demultiplexing and before the mapping, the reads were trimmed to remove the adaptor sequences from the first amplification. FASTQ files were mapped to the canFam4 reference using HISAT2 and counted by featureCounts from the Subread R package. Normalization was performed with R package DESeq2 v1.24.0^43^. Flags were computed from counts normalized to the mean coverage. All normalized counts <20 were considered as background (flag 0) and >=20 as signal (flag=1). P50 lists used for the statistical analysis regroup the genes showing flag=1 for at least half of the compared samples. For differential gene expression analysis, a 2-fold fold change threshold was applied as well as a Bonferroni correction with threshold for significance set as adjusted P-values < 0.05.For reanalysis of transcriptomic data from obstructed mouse kidneys^44^, we downloaded from the Gene Expression Omnibus database (https://www.ncbi.nlm.nih.gov/geo/) the expression matrix of mRNAs expressions of 7-day-UUO kidneys of under the accession number GSE145053.

### Human-iPSC derived kidney organoids

Human induced pluripotent stem cells were maintained on Vitronectin-coated surfaces in mTeSR Plus medium (StemCell Technologies, #100-0276). One day prior to differentiation, cells were dissociated using ReLeSR (StemCell Technologies, #100-0483), pelleted by centrifugation (300 × g, 5 min), and replated at a density of 5 × 10⁵ viable cells per well in 6-well plates (Corning, #3506) pre-coated with growth factor-reduced Matrigel (Corning, #354277; 1:100 dilution in DMEM/F-12, Gibco, #31330038; 30 min at room temperature) in mTeSR Plus medium. On Day 1, culture medium was replaced with APEL medium (StemCell Technologies, #05270) supplemented with 8 µM CHIR99021 (Bio-Techne, #4423/10) to initiate mesoderm induction, and cells were cultured for 3 days. On Day 4, medium was exchanged for APEL containing 200 ng/mL FGF9 (Bio-Techne, #273-F9-025) and 1 µg/mL heparin (StemCell Technologies, #07980), refreshed every 2 days until Day 8. On Day 8, cells were washed with D-PBS (Gibco, #14190169), dissociated with Cell Dissociation Buffer PBS-based (Gibco, #13151014; 8 min, 37°C), and the reaction was stopped in KnockOut DMEM (Gibco, #10829018) supplemented with 1% BSA (Euromedex, #04-100-812-C). Cells were pelleted (300 × g, 5 min), resuspended in APEL containing 5 µM CHIR99021, and counted by Trypan Blue exclusion. Pellets of 3 × 10⁵ cells were formed by centrifugation and transferred onto Transwell filter inserts (Corning, #3450) — three pellets per insert for 6-well formats, one per insert for 12-well formats — in 1.2 mL or 400 µL APEL/5 µM CHIR99021 per well, respectively. After 1 hour at 37°C, medium was replaced with APEL containing 200 ng/mL FGF9, 1 µg/mL heparin, and 1x antibiotic-antimycotic (Gibco, #15240062), refreshed every 2 days through day 14. From day 14, organoids were cultured in APEL supplemented with 1 µg/mL heparin and 1x antibiotic-antimycotic alone, with medium changes every 2 days until day 22, at which point organoids were considered fully differentiated.

### Single-cell transcriptomic analysis of ciliary and NF-κB signatures in the Kramann kidney atlas

Single-cell RNA sequencing data from the human kidney atlas^33^ were analyzed using the Seurat R package (v4). The pre-processed Seurat object was loaded with cell type annotations as previously described^45^, encompassing 34 cell populations across tubular, stromal, endothelial, and immune compartments.

#### Gene signatures

The ciliary signature was derived from the Syscilia ciliome database^46^ and comprised all genes with annotated ciliary localization. The NF-κB pathway signature corresponded to the BIOCARTA_NFKB_PATHWAY gene set. For each signature, only genes present in the dataset’s feature space were retained.

#### Signature scoring

Gene signature scores were computed using Seurat (v4) in R. Log-normalized expression values were extracted from the Seurat object data slot using GetAssayData(). For each signature, only genes present in the dataset were retained. Expression values were then z-score normalized per gene across all cells (subtracting the gene-wise mean and dividing by the standard deviation). The per-cell signature score was defined as the mean z-score across all retained signature genes. Scores were projected onto UMAP embeddings and visualized using FeaturePlot(), with a gray-to-red color scale and quantile-based cutoffs (10th–90th percentiles), and as violin plots grouped by cell annotation using VlnPlot().

#### Correlation analysis

Spearman correlation coefficients (ρ) between ciliary and NF-κB signature scores were computed independently for each annotated cell type, and p-values were adjusted for multiple testing using the Benjamini-Hochberg method. Results were visualized as scatter plots with per-cell-type ρ and FDR annotations. Expression of HNF4A and SOX9 was visualized by violin plot across tubular cell types.

### Data availability

The transcriptomic data supporting the findings of this study are openly available in the public domain. The deposition is ongoing. An accession name will be provided with the first revised version of the article. Additional RNAseq datasets were derived from the following resources available in the public domain: https://www.ncbi.nlm.nih.gov/geo/ (GSE145053). All the mice and cell models used in this study can be obtained upon request to the corresponding author, with the agreement of the scientist who generated the initial transgenic line.

### Statistical Analysis

Data were expressed as means. Shapiro-Wilk test was performed to verify the distribution of the data. For data that did not follow a normal distribution, differences between groups were evaluated using Student’s Mann-Whitney test (when only two groups were compared) or Kruskal-Wallis test with Dunn’s multiple comparison test (when testing more comparisons). For data that followed normal distribution, Student’s unpaired *t* test with Welch’s correction was used when comparing only two sets of data. When testing more than 2 groups, One-way ANOVA with Tukey’s multiple comparison test or ANOVA Brown-Forsythe test with Tamhane’s T2 multiple comparison test were used depending on variance differences. To determine if there is a significant difference between two proportions (example ciliogenesis), we used Fisher’s exact test. To determine if there is a significant difference between more than two proportions, we used a χ2 test. P < 0.05 was considered statistically significant. The statistical analysis was performed using GraphPad Prism V10 software. All image analyses and mouse phenotypic analyses were performed in a blinded fashion.

Data are expressed as mean ± standard error of the mean (SEM). Statistical analyses were performed on the data generated from this study using GraphPad Prism 9.2.0 for Windows (GraphPad Software, Inc.) and are depicted in Table 2 below. Statistical significance was set at p < 0.05. In multi-dose cell assays, DF-003 activity was calculated via logarithmic interpolation. Concentration-response curves were fitted with a four-parameter logistic nonlinear regression model and the IC_50_ values calculated by GraphPad Prism. Observations which were beyond the mean of each group by 3 standard deviations (SD) were considered outliers but were not excluded from graphs or statistical analyses.

### Study Approval

All animal experiments were conducted according to the guidelines of the National Institutes of Health Guide for the Care and Use of Laboratory Animals, and were approved by regional authorities (the French “Ministère de l’Enseignement, de la Recherche et de l’Innovation” under protocol numbers 32143-2021042212059917 and 39771-2022081016482946 and the Institutional Animal Care and Use Committees (IACUC) at Shanghai Yao Yuan Biotechnology Ltd., Zhejiang Yao Yuan Biotechnology Ltd. and Oujiang Laboratory under protocol number DFSH00320220910-001).

## ACKNOWLEDGMENTS

We thank the members of the LEAT, histology, genomics, bioinformatics, cell imaging and image analysis facilities (S.F.R Necker INSERM US24, Paris, France) and of the mouse renal physiology facility (Cordeliers Research Center, Paris, France) for technical assistance. Emmanuelle Bille, Matthieu Coureuil, Ines Imbit, Catharina Svanborg, Keira Melican and Agneta Richter Dahlfors for providing bacterial strains and thoughtful discussion.

We are grateful for support to the Ultrastructural BioImaging Core Facility equipment from the GIS-IBISA, the French Government Programme Investissements d’Avenir France BioImaging (FBI, N° ANR-10-INSB-04-01 / ANR-24-INBS-0005FBI BIOGEN), the DIM1 Health and the French government (Agence Nationale de la Recherche) Investissement d’Avenir programme, Laboratoire d’Excellence “Integrative Biology of Emerging Infectious Diseases” (ANR-10-LABX-62-IBEID).

JM received a PhD grant from the Ministère de l’Education Nationale, de la Recherche et de la Technologie (MENRT). The team of JE was supported for this work by the ANR (grants PureMagRupture, DynamicHostPathOMICs, and RabReprogram), and they are member of the LabEx “IBEID” and “Milieu Interieur”. MAI received support from the Agence Nationale de la Recherche (ANR, ANR-22-CE15-0022-01 ; ANR-23-CE14-0025-01). AV received support from the Fondation pour la recherche Médicale (FRM, grant reference: ARF20150934110), from the Agence Nationale de la Recherche (ANR, grant reference: ANR-23-CE14-0025-01) and the program “*Investissement d’Avenir*” launched by the French Government and implemented by ANR, with the reference « ANR-18-IdEx-0001 » as part of its program « *Emergence* ». FB was supported by the Agence Nationale de la Recherche (ANR, grant references: ANR-19-CE14-0016; ANR-23-CE14-0025-03); the program “*Investissement d’Avenir*” launched by the French Government and implemented by ANR, with the reference « ANR-18-IdEx-0001 » as part of its program « *Emergence* », the SFNDT (*Société Française de Néphrologie, Dialyse et Transplantation*: D24025KK), the AIRG (*Association pour l’information et la recherche sur les maladies génétiques rénales*).

## AUTHOR CONTRIBUTIONS

Design of the work: MR, JF, HL, AV and FB

Acquisition, analysis and interpretation of data: AA, JM, FK, TB, SC, CC, A-S.C, MMN, NC, NG, MGT, MQ, CX, LA, MR, CA, JF, MAI, AV, FB

Drafted and revised the manuscript: CC, E.W.K, JE, MR, CA, JF, SS, FT, MAI, HL, AV, FB All authors approved the final version of the manuscript.

