## Extended Data tables and Figures for "A cilia-dependent inflammatory programme links bacterial detection to kidney disease"

### Supplementary Data

### TABLE OF CONTENTS

**Supplementary Tables**

**Supplementary Table 1**: Antibodies list

**Supplementary Table 2**: Primer pairs used for qRT-PCR.

**Supplementary Table 3**: Bacterial strains used.

**Supplementary Table 2**: Reagents.

**Supplementary Figure Legends**

**Extended Data Fig. 1**

**Extended Data Fig. 2**

**Extended Data Fig. 3**

**Extended Data Fig. 4**

**Extended Data Fig. 5**

**Extended Data Fig. 6**

**Extended Data Fig. 7**

**Extended Data Fig. 8**

**Extended Data Fig. 9**

**Extended Data Fig. 10**

**Extended Data Fig. 11**

**Extended Data Fig. 12**

**Supplementary Table 1: Antibodies used for immunohistochemistry, immunofluorescence or western blotting**

| Antibody | Company | Reference | Specie | FFPE Antigen retrieval |
|---|---|---|---|---|
| Acetylated-Tubulin | Sigma | T-6793 | Mouse | None/Citrate/TE |
| Acetylated-Tubulin A488 | Santa Cruz | sc-23950 AF488 | Mouse | None/Citrate/TE |
| Acetylated-Tubulin A647 | Santa Cruz | sc-23950 AF 555 | Mouse | None/Citrate/TE |
| Alpha-Tubulin | Sigma | T5168 | mouse | - |
| ANKS6 | Sigma | HPA008355-100 | Rabbit | TE |
| AQP2 | Sigma | A7310 | Rabbit | None/Citrate/TE |
| AQP2 | Santa Cruz | sc-515770 | Mouse | None/Citrate/TE |
| ARL13B | Proteintech | 17711-1-AP | Rabbit | TE |
| CALB1 | Santa Cruz | sc-365360 | mouse | Citrate/TE |
| Ecadherin | Life technologies | 131900 | rat | None |
| F4/80 | Cell Signaling Technology | 70076 | Rabbit | PK |
| Gamma-Tubulin | Sigma | T6557 | Mouse | TE |
| GFP | Alves | GFP-1010 | Chicken | TE |
| IFT88 | ProteinTech | 13967-1-AP | Rabbit | - |
| KIF3A | BD Transduction Laboratories™ | 611508 | Mouse | - |
| LPS antibody [2D7/1] | Abcam | ab35654 | Mouse | |
| LY6B.2 | Abcam | ab53457 | Rat | None/Citrate/TE |
| UMOD | AbDSerotec/BioRad | 8595-0054 | Sheep | TE |
| WGA Fluorescein | Vector Laboratories | FL-1021 | NA | None/Citrate/TE |
| WGA Rhodamine | Vector Laboratories | RL-1023 | NA | None/Citrate/TE |

PK : incubate section in proteinase K 1 µL (Roche #03 115 887 001, 18+/-4 mg/mL) 1:1000 in 50mM Tris 1mM EDTA pH 8 for 2 minutes
TE : heat in TE buffer (10mM Tris Base, 1mM EDTA Solution, 0.05% Tween 20, pH 9) 15 min at 120°C high pressure in a pressure cooker (TintoTertriver; BioSB)
Citrate : heat in 1X citrate buffer (Zytomed Systems, ZUC028) 15 min at 120°C at high pressure in a pressure in a pressure cooker (TintoTertriver; BioSB)

| Antibody | Company | Reference |
|---|---|---|
| Donkey anti-mouse A488 | Life Technologies | A21202 |
| Donkey anti-mouse A555 | Life Technologies | A31570 |
| Donkey anti-mouse A647 | Life Technologies | A31571 |
| Goat anti-IgG1 mouse A647 | Life Technologies | A21240 |
| Goat anti-IgG2b mouse A555 | Life Technologies | A21147 |
| Goat anti-IgG2b mouse A488 | Life Technologies | A21141 |
| Goat anti-IgG2a mouse A488 | Life Technologies | A21131 |
| Mouse HRP | GE Healthcare | NA931-1ML |
| Rabbit HRP | GE Healthcare | NA934V-1ML |
| Donkey anti-rabbit A488 | Life Technologies | A21206 |
| Donkey anti-rabbit A555 | Life Technologies | A31572 |
| Donkey anti-rabbit A647 | Life Technologies | A31573 |
| Donkey anti-goat A488 | Life Technologies | A11055 |
| Donkey anti-goat A555 | Life Technologies | A21432 |
| Donkey anti-goat A647 | Life Technologies | A21447 |
| Goat anti-rat HRP | Vector | PI-9400 |
| Goat anti-chicken A488 | Life Technologies | A11039 |
| anti-streptavidin A488 | Life Technologies | S32354 |
| anti-streptavidin A555 | ThermoFischer | S32355 |
| anti-streptavidin A647 | ThermoFischer | S32357 |

| | | |
|---|---|---|
| Hoechst 33342 | Molecular Probes | H1399 |

**Supplementary Table 2: Primer pairs used for qRT-PCR.**

| Gene name | Specie | Forward Primer (5' to 3') | Reverse Primer (5' to 3') |
|---|---|---|---|
| *Acta2* | Mouse | CATGCGTCTGGACTTGGCTG | GACAATCTCACGCTCGGCAGTAG |
| *Aqp2* | Mouse | TAGCCCTGCTCTCTCCATTGGTTT | AAACTTGCCAGTGACAACTGCTGG |
| *Ccl2* | Mouse | AGTAGGCTGGAGAGCTACAA | GTATGTCTGGACCCATTCCTTC |
| *Ccl5* | Mouse | CCAATCTTGCAGTCGTGTTTG | ACCCTCTATCCTAGCTCATCTC |
| *Col1a1* | Mouse | GCCGCAAAGAGTCTACATGTCTAG | TGGCAGATACAGATCAAGCATACC |
| *Col3a1* | Mouse | GGACCAGCAGGAACTAATGGTAT | GTTCTCCAGGTGATCCATCTTT |
| *Col4a1* | Mouse | GTCTGGCTTCTGCTGCTCTTC | CCTTCACGCCATGACAGTCA |
| *Cx3cl1* | Mouse | GCTTTGCTCATCCGCTATCA | GTCTTGGACCCATTTCTCCTTC |
| *Cxcl1* | Mouse | CGAAGTCATAGCCACACTCAA | GAGCAGTCTGTCTTCTTTCTCC |
| *Cxcl10* | Mouse | GGCCATAGGGAAGCTTGAAA | CAGACATCTCTGCTCATCATTCT |
| *Cxcl16* | Mouse | ATCAGGTTCCAGTTGCAGTC | CATGACCAGTTCCACACTCTT |
| *Cxcl17* | Mouse | CCTTCCTTCTGTTGCTTCCA | TTCCAAGAGCCACCTCCTA |
| *Hnf4a* | Mouse | TGAGGAAGAACCACATGTACTC | CGACACTGGTTCCTCTTATCTT |
| *Hprt* | Mouse | GTTAAGCAGTACAGCCCCAAA | AGGGCATATCCAACAACAAACTT |
| *Ift20* | Mouse | GGTGGTCTAATTGAGCTTGTTG | CTCTGCTTCGCTATGGACTT |
| *Il1rn* | Mouse | TTGTGCCAAGTCTGGAGATG | CTCAGAGCGGATGAAGGTAAAG |
| *Il33* | Mouse | TGCCTCCCTGAGTACATACA | CTGGTCTTGCTCTTGGTCTTT |
| *Il34* | Mouse | GATATGGACTCTGACCCAAGATAAG | AGCAATCCTGTAGTTGATGGG |
| *Kif3a* | Mouse | GACATAGGAGAATGGCAGCTAAA | CGACTTCAAATGGGTCTCTCTC |
| *Lcn2* | Mouse | GGACCAGGGCTGTCGCTACT | GGTGGCCACTTGCACATTGT |
| *Lgals9* | Mouse | CAGATGCTACGAGGTTCCATATC | GTGTTTCGGACAACAGCATTC |
| *Lrp2* | Mouse | CCAGCAGAGGCGTGATTG | TAAAGGAAGGAGCCGTGGAC |
| *Pdgfb* | Mouse | GGCAAGCACCGAAAGTTTAAG | TAAATAACCCTGCCCACACTC |
| *Ppia* | Mouse | GGCTATAAGGGTTCCTCCTTTC | TTTCTCTCCGTAGATGGACCT |
| *Rpl13* | Mouse | GCTCCAAGCTCATCCTGTT | GGTGGCCAGCTTAAGTTCT |
| *Scnn1a* | Mouse | AGTGTGGCTGTGCCTACATCTTCT | TTGAGTAGCCAGCAGAGAGCTTGT |
| *Slc26a1* | Mouse | TGATGGCCGGGCTTTATCAG | TCGAGCAGTGGTTGTGAGAG |
| *Slc34a1* | Mouse | GGCTTGTGGTTGCTTCCTTCAACA | TGCTGGTGATCACAGACTTGTCCA |
| *Tgfb1* | Mouse | GGGAAGCAGTGCCCGAACCC | TGGGGGTCAGCAGCCGGTTA |

| Gene name | Specie | Forward Primer (5' to 3') | Reverse Primer (5' to 3') |
|---|---|---|---|
| *Ccl2* | Dog | CTGCTGCTATACACTCACCAATA | TCAGCACAGATCTCCTTGTTTAG |
| *Cxcl8* | Dog | CTCTCTGTGAAGCTGCAGTTCTG | GGAAAGGTGTGGAGTGTGTTTTT |
| *Cxcl10* | Dog | TGAAATGATTCCTGCAAGTCCAT | TCTCCCCACTCTTTTTCATTGTG |
| *Gapdh* | Dog | CATGTTTGTGATGGGCGTGAACCA | TTTGGCTAGAGGAGCCAAGCAGTT |
| *Hnf4a* | Dog | CATGTACTCCTGCAGGTTTAGTC | GCCCGGAAGCACTTCTTTA |
| *Hprt* | Dog | TGCTCGAGATGTGATGAAGG | TCCCCTGTTGACTGGTCATT |
| *Pdgfb* | Dog | GCTGCAACAACCGCAAC | TGGCTTCTTCCGTACAATCTC |
| *Sdha* | Dog | TGGGTCCATCCATCGCATA | GAAGTAGGTGCGCCCATAACC |
| *Wnt5a* | Dog | GCAGCACCGTGGATAACA | CACCGCGTACGTGAAGG |

| Gene name | Specie | Forward Primer (5' to 3') | Reverse Primer (5' to 3') |
|---|---|---|---|
| *CCL2* | human | TCATAGCAGCCACCTTCATTC | CTCTGCACTGAGATCTTCCTATTG |
| *CXCL1* | human | ACTCAAGAATGGGCGGAAAG | CCCTTCTGGTCAGTTGGATTT |
| *CXCL8* | human | GTGCATAAAGACATACTCCAAACC | CAGAGCTCTCTTCCATCAGAAA |
| *CXCL10* | human | CCATTCTGATTTGCTGCCTTATC | TACTAATGCTGATGCAGGTACAG |
| *PPIA* | human | GGTCCCAAAGACAGCAGAAA | GGTCCCAAAGACAGCAGAAA |
| *SDHA* | human | GCCGTGGTCGAGCTAGAAAA | CACGCTGATAAATCTTCCCATCT |
| *TBP* | human | TCCACAGTGAATCTTGGTTGT | TCCACAGTGAATCTTGGTTGT |

| Gene name | Specie | Forward Primer (5' to 3') | Reverse Primer (5' to 3') |
|---|---|---|---|
| *Acta2* | rat | AGGAGCATCCGACCTTGCTA | AGAGTCCAGCACAATACCAGTT |
| *Ccl2* | rat | TGTAGTTCTCCAGCCGACTC | GCCTGTTGTTCACAGTTGCT |
| *Col1a1* | rat | ATCGACCCTAACCAAGGCTG | CGCTTCCATACTCGAACTGG |
| *Cxcl1* | rat | GCCACACTCAAGAATGGTCG | GACGCCATCGGTGCAATCTA |
| *Cxcl10* | rat | GAATCCGGAATCTGAGGCCA | CGTCTCTCTGCTGTCCATCG |
| *Tgfb* | rat | GGACCGCAACAACGCAATCT | AGAGTTCTACGTGTTGCTCCA |
| *Ubc* | rat | ACACCAAGAAGGTCAAACAGGAA | CCCAAGAACAAGCACAAGAAGG |

| Gene name | Specie | Forward Primer (5' to 3') | Reverse Primer (5' to 3') |
|---|---|---|---|
| *CCL2* | human | AGAATCACCAGCAGCAAGTGTCC | TCCTGAACCCACTTCTGCTTGG |
| *CCL5* | human | CCTGCTGCTTTGCCTACATTGC | ACACACTTGGCGGTTCTTTCGG |
| *CXCL10* | human | GGTGAGAAGAGATGTCTGAATCC | GTCCATCCTTGGAAGCACTGCA |
| *CXCL2* | human | GGCAGAAAGCTTGTCTCAACCC | CTCCTTCAGGAACAGCCACCAA |
| *CXCL8* | human | GAGAGTGATTGAGAGTGGACCAC | CACAACCCTCTGCACCCAGTTT |
| *GAPDH* | human | AATTCCATGGCACCGTCAAG | TGGACTCCACGACGTACTCA |
| *IL1b* | human | CAGAAGTACCTGAGCTCGCC | AGATTCGTAGCTGGATGCCG |
| *IL6* | human | AGTGAGGAACAAGCCAGAGC | AGCTGCGCAGAATGAGATGA |
| *TNFa* | human | CTCTTCTGCCTGCTGCACTTTG | ATGGGCTACAGGCTTGTCACTC |

**Supplementary Table 3: Bacterial strains used**

| Strain | Original strain | Gift from | Original publication / origin |
|---|---|---|---|
| UTI89 | UTI89 | Molly Ingersoll | PMID: 26182347 |
| UTI89-RFP-kan$^R$ | UTI89 | Molly Ingersoll | PMID: 26182347 |
| UTI89-GFP-amp | UTI89 | Molly Ingersoll | PMID: 26182347 |
| CFT073 | CFT073 | Molly Ingersoll from ? | PMID: 26182347 |
| ARD42 W3110 cobS::W(PLtetO-1-gfp+), cmr | E.Coli K12 lab strain | Agneta Richter Dahlfors | PMID: 21383970 |
| 83972 | 83972 | Catharina Svanborg/Inès Ambite | PMID: 31181116 |
| 83972+FIM | 83972 | Catharina Svanborg/Inès Ambite | PMID: 31181116 |
| 83972+PAP | 83972 | Catharina Svanborg/Inès Ambite | PMID: 31181116 |
| UTI89 Parental | UTI89 | Cécile Arriemerlou | PMID: 26182347 |
| AIEC LF82 ∆FLA | AIECLF82 | David Skurnik | PMID: 12694621 |

| | | | |
|---|---|---|---|
| Proteus mirabillis (CIP 103 181T) | NA | Emmanuelle Bille | Biological Resource Center of Institut Pasteur |
| Pseudomonas aeruginosa (CIP 105 149) | NA | Emmanuelle Bille | Biological Resource Center of Institut Pasteur |
| Pseudomonas aeruginosa (CIP 76 110) | NA | Emmanuelle Bille | Biological Resource Center of Institut Pasteur |
| Klebsiella pneumoniae (CIP 82.91T) | NA | Emmanuelle Bille | Biological Resource Center of Institut Pasteur |
| Enterobacter cloacae (CIP 60.85T) | NA | Emmanuelle Bille | Biological Resource Center of Institut Pasteur |
| Enterococcus faecalis 72.27 (CIP 103 907) | NA | Emmanuelle Bille | Biological Resource Center of Institut Pasteur |
| Enterococcus faecalis OG1RF (CIP 103 907) | NA | Molly Ingersoll | PMID: 28893918 |
| Neisseria elongata (CIP 72.27) | NA | Emmanuelle Bille | Biological Resource Center of Institut Pasteur |

**Supplementary Table 4: Reagents**

| Reagent ID | Reagent full name | Reference | Comments |
|---|---|---|---|
| **ADPh** | ADP-L-glycero-β-D-manno-heptose | tlrl-adph-l, InvivoGen | Dissolved in endotoxin-free water at 1mg/mL and used at 10µg/mL |
| **CLI-095** | CLI-095 - TLR4 Signaling Inhibitor | tlrl-cli95-4, InvivoGen | Dissolved in DMSO at 2.5 mM and used at 5µM |
| **DF-003** | DF-003 | PMID: 40925900 | Synthesized by Shanghai Yao Yuan Biotechnology Co., Ltd. |
| **DF-006** | D-glycero-D-manno-6-fluoro-heptose-1β- S-ADP | PMID: 35699669 | Manufactured by Drug Farm Inc., Lot CKo125064-01-pxy-0456-02 |
| **FLA** | Flagellin, FLA-ST Ultrapure | tlrl-epstfla, InvivoGen | Dissolved in steril water at 100 µg/mL and used at 100ng/mL |
| **LPS** | Lipopolysaccharide from Escherichia coli O55:B5 | tlrl-b5lps, Invivogen | Dissolved in endotoxin-free at 5 mg/ml and use at 5µg/mL |
| **MDP** | Muramyl dipeptide | tlrl-mdp, InvivoGen | Dissolved in steril water at 10 mg/mL and used at 10µg/mL |
| **Pam2** | Synthetic diacylated lipopeptide, Pam2CSK4 | tlrl-pm2s-1, InvivoGen | Dissolved in endotoxin-free at 5 mg/ml and use at 5µg/mL |
| **Pam3** | Synthetic triacylated lipoprotein, Pam3CSK4 | tlrl-pms, InvivoGen | Dissolved in endotoxin-free at 5 mg/ml and use at 5µg/mL |
| **TAK1-i** | Takinib | 6430, Tocris Bioscience | Dissolved in DMSO at 30mg/mL and used at 3.2 µg/mL |
| **TBK1-i** | Amlexanox | inh-amx, InvivoGen | Dissolved in DMSO at 30mg/mL and used at 7.5µg/mL |
| **Tri-DAP** | L-Ala-g-D-Glu-mDAP | tlrl-tdap, InvivoGen | Dissolved in steril water at 10 mg/mL and used at 10µg/mL |

Extended Data Fig. 1

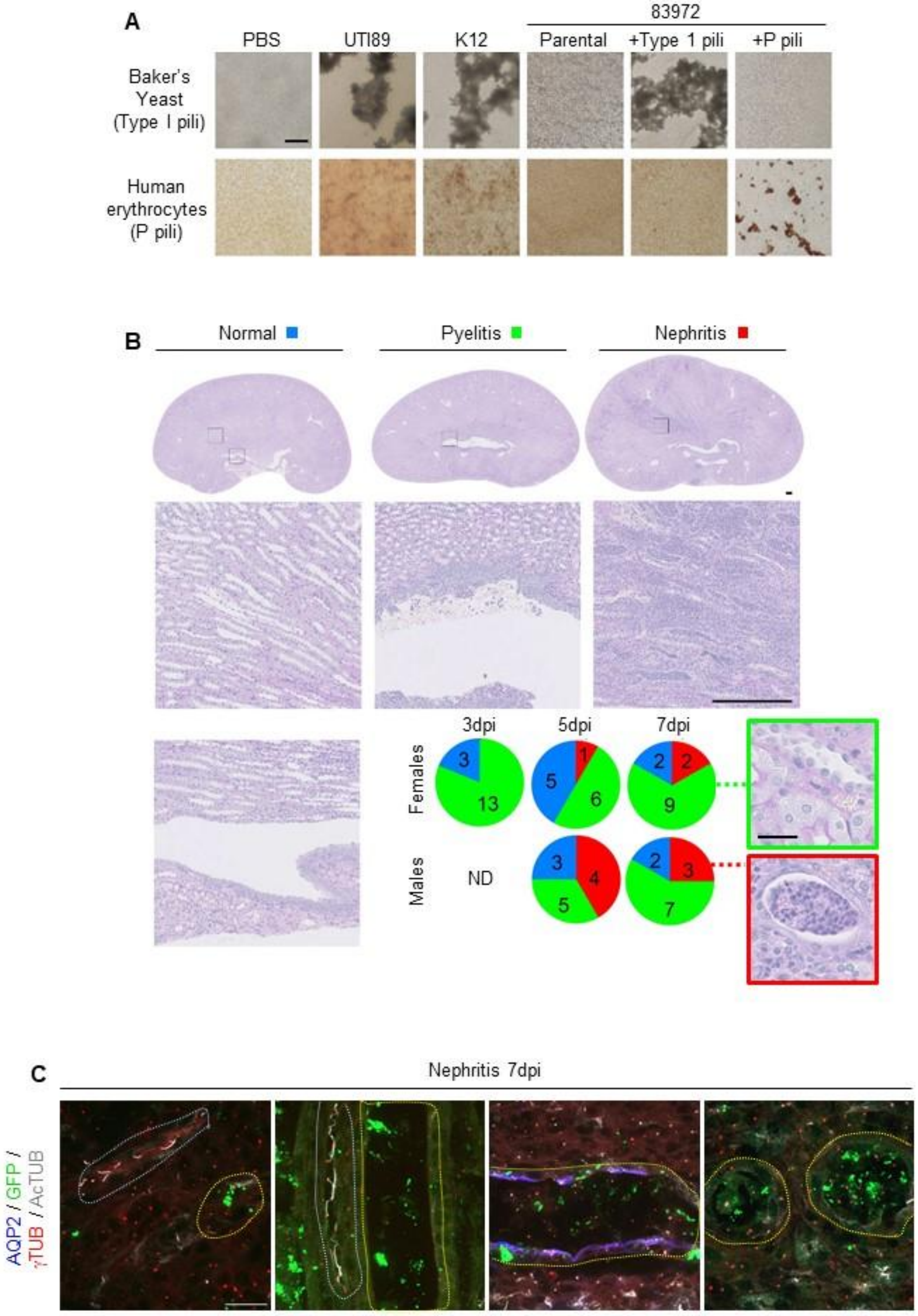

**Extended Data Figure 1.**

**(A)** Representative images of baker's yeast or human erythrocyte agglutination by the indicated UPEC strains. Agglutination of baker's yeast and human erythrocytes indicates functional type 1 pili and P pili, respectively. Scale bar: 200 µm. **(B)** Representative PAS-stained kidney sections from 8-week-old C3H/HeN mice after bladder instillation of UPEC (UTI89), showing normal kidneys, isolated pyelitis, or overt nephritis. Pie charts show the distribution of male and female mice within each histological severity class, 3, 5, or 7 days after instillation. Numbers within pies indicate the number of mice. Insert in green and red shows examples of normal tubules in kidneys with pyelitis (green) or infected tubules with flattened epithelium (red) 7 dpi. Scale bars: 250 µm (20µm for inserts). **(C)** Representative images of kidney sections from C3H mice 7 days after bladder instillation of GFP-labelled UTI89, immunostained with antibodies against aquaporin 2 (AQP2) and acetylated α-tubulin (AcTUB), showing deciliation of infected tubules (yellow dashed area) contrasting with ciliated, uninfected tubules displaying long primary cilia (light blue dashed area). Scale bar: 20 µm.

Extended Data Fig. 2

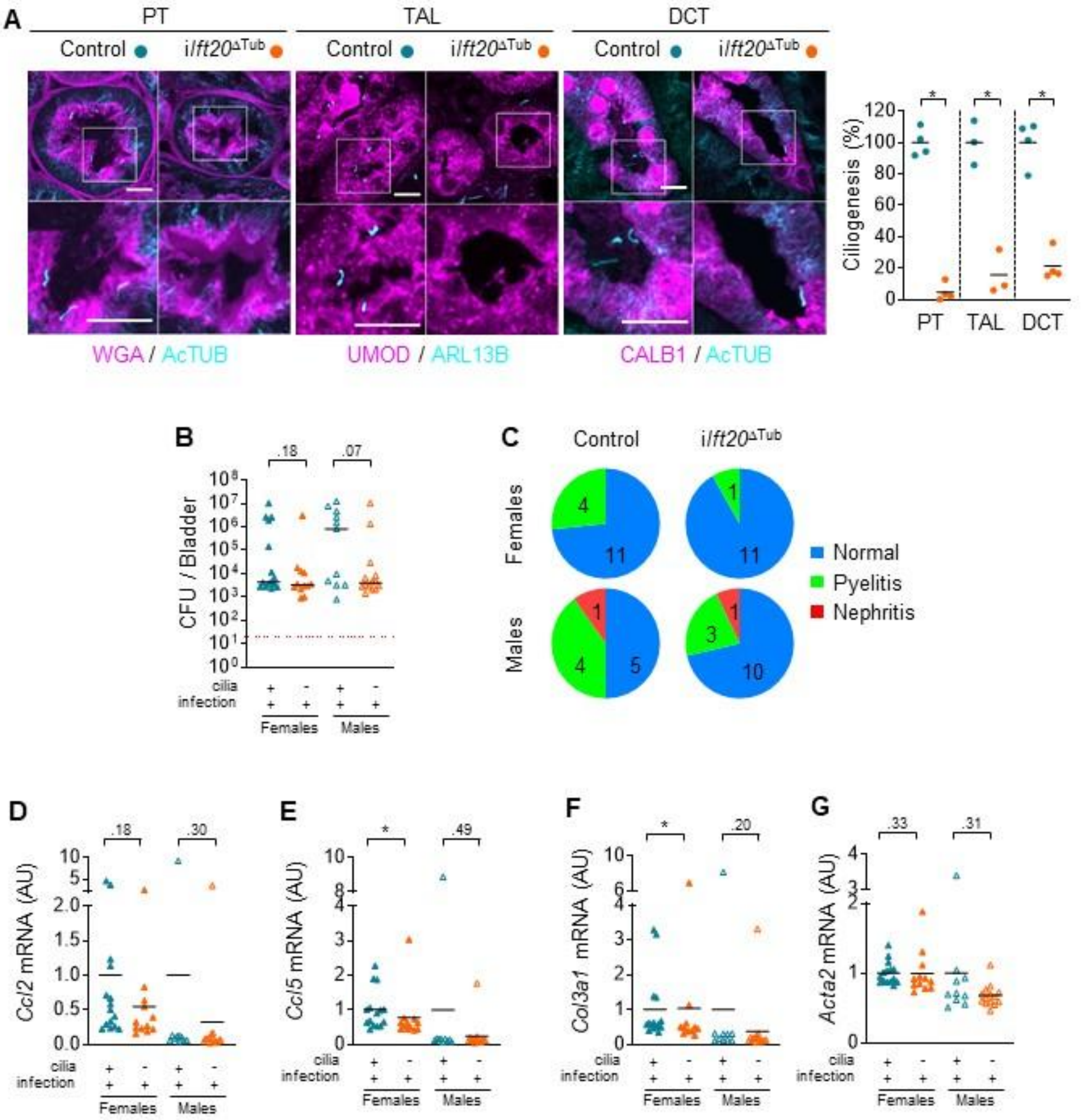

**Extended Data Figure 2.**

**(A)** Representative images and quantification of primary cilia (AcTUB+ or ARL13B+, blue) in proximal tubules (PT, with WGA+ brush border), thick ascending loop of Henle (TAL, UMOD+) and distal convoluted tubules (DCT, CALB1+) of kidneys from 8-week-old control and *Ift20*$^{\Delta Tub}$ mice, 2 weeks after the completion of doxycycline treatment. Scale bar: 10µm. Bars indicate mean. Each dot represents a female mouse. Mann-Whitney t-test: *P<0.05. AU: arbitrary units. **(B)** Colony forming units (CFU) in the bladder of 11-week-old control and *Ift20*$^{\Delta Tub}$ mice, 7 days after intravesical instillation of UPEC strain UTI89. **(C)** Pie charts showing the distribution of male and female mice within each histological severity class 7 dpi. Numbers within pies indicate the number of mice. Colors represent normal tubules in kidneys (blue), with pyelitis (green) or infected tubules with flattened epithelium (red) 7 dpi. **(D-G)** qPCR quantification of *Ccl2* (D), *Ccl5* (E), *Col3a1* (F) and *Acta2* (G) mRNA in infected kidneys from 11-week-old control and *Ift20*$^{\Delta Tub}$ mice, 7 days after intravesical instillation of UTI89. **(B, D-E)** Bars indicate mean. Each dot represents a mouse. Filled symbols represent females while empty symbols represent males. Mann-Whitney t-test: *P<0.05. AU: arbitrary units.

Extended Data Fig. 3

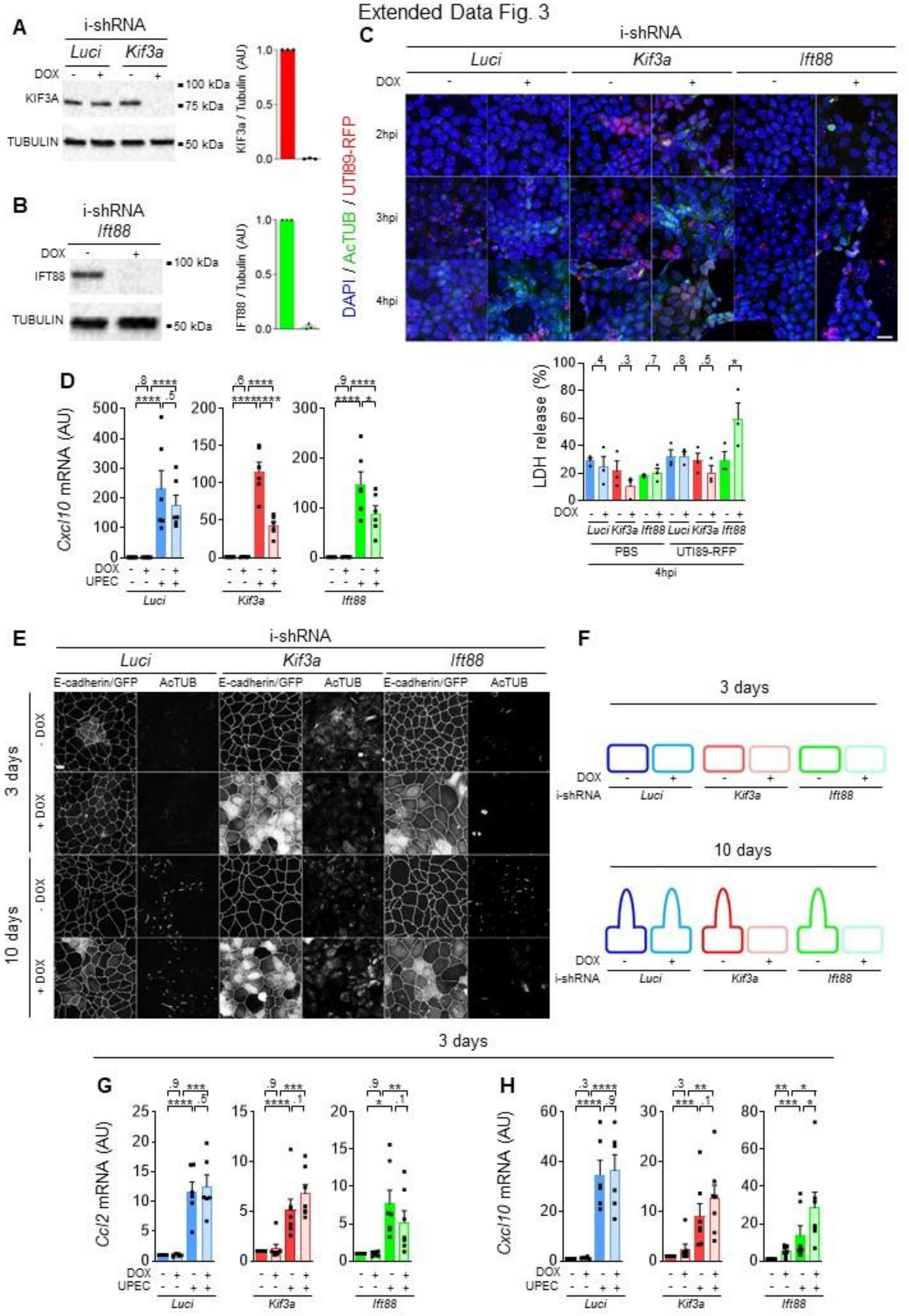

**Extended Data Figure 3.**

**(A–B)** Representative western blots and quantification of KIF3A (A) and IFT88 (B) in kidney cells (MDCK) expressing doxycycline inducible shRNA targeting *Kif3a* (red, A) or *Ift88* (green, B) after 2 days of culture. **(C)** Representative images of primary cilia (AcTUB+, green), living UPEC (UTI89.RFP, red) and DNA (blue) in 10-day-cultured shRNA-inducible MDCK epithelial layers, 2, 3, or 4 hours following UPEC infection, and quantification of LDH release, 4 hours post UPEC infection (hpi). Scale bar 20μm. Ratio paired t-test: *P<0.05. **(D)** qPCR quantification of *Cxcl10* mRNA at 10 days of culture in the same cells treated for 6 hours by heat killed UPEC (UTI89) or vehicle (PBS). Paired one-way RM ANOVA (log-normal) with Geisser Greenhouse correction followed by Tukey test: *P<0.05, ***P<0.001, ****P<0.00001. Each dot per condition represents an independent experiment. AU: arbitrary units. **(E-F)** Representative labelling of E-cadherin and primary cilia (AcTUB+) in the same cell line treated or not with doxycycline for 3 or 10 days after seeding. Doxycycline also induces the expression of green fluorescent protein (GFP), which is visible in the E-cadherin channel. Most cells lack cilia 3 days after seeding (E). Scale bar 10μm. **(G-H)** qPCR quantification of *Ccl2* (G) and *Cxcl10* (H) mRNA after 3 days of culture in the same cell lines treated for 6 hours with heat-killed UPEC (UTI89) or vehicle (PBS). Each dot represents an independent experiment. Paired one-way RM ANOVA (log-normal) with Geisser Greenhouse correction followed by Tukey test: *P < 0.05, **P < 0.01, ***P < 0.001, ****P<0.00001. AU: arbitrary units.

Extended Data Fig. 4

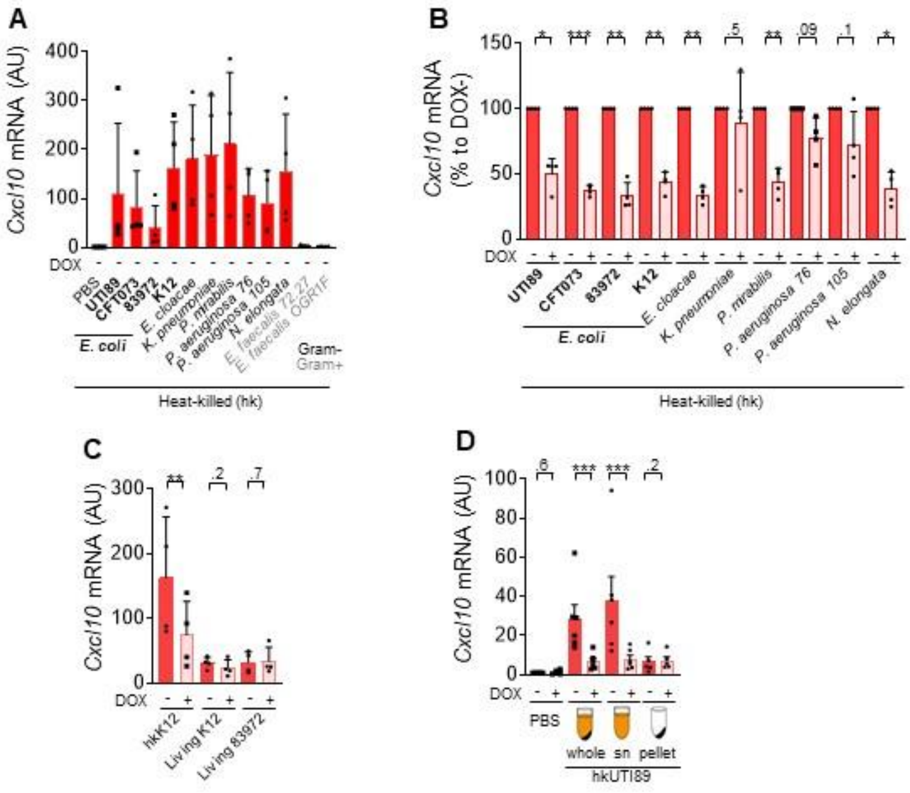

**Extended Data Figure 4.**

**(A-B)** qPCR quantification of *Cxcl10* mRNA in 10 days cultured *Kif3a* i-shRNA MDCK cells treated for 6 hours by the indicated heat killed Gram-negative (black) or -positive (grey) bacteria. Absolute induction in ciliated cells and relative effect of cilia ablation are shown in A and B, respectively. **(C)** qPCR quantification of *Cxcl10* mRNA in *Kif3a* i-shRNA MDCK cells treated for 6 hours by UPEC heat-killed (hkK12) or not (living K12 and living 83972). **(D)** qPCR quantification of *Cxcl10* mRNA in *Kif3a* i-shRNA MDCK cells treated for 6 hours by UPEC heat-killed (hkUTI89) in PBS (whole) and the corresponding supernatant (sn) or pellet fractions of the preparation. **(A-D)** Ratio paired *t* test: *P<0.05, **P<0.01, ***P<0.001. Each dot represents an experimental n. AU: arbitrary units.

Extended Data Fig. 5

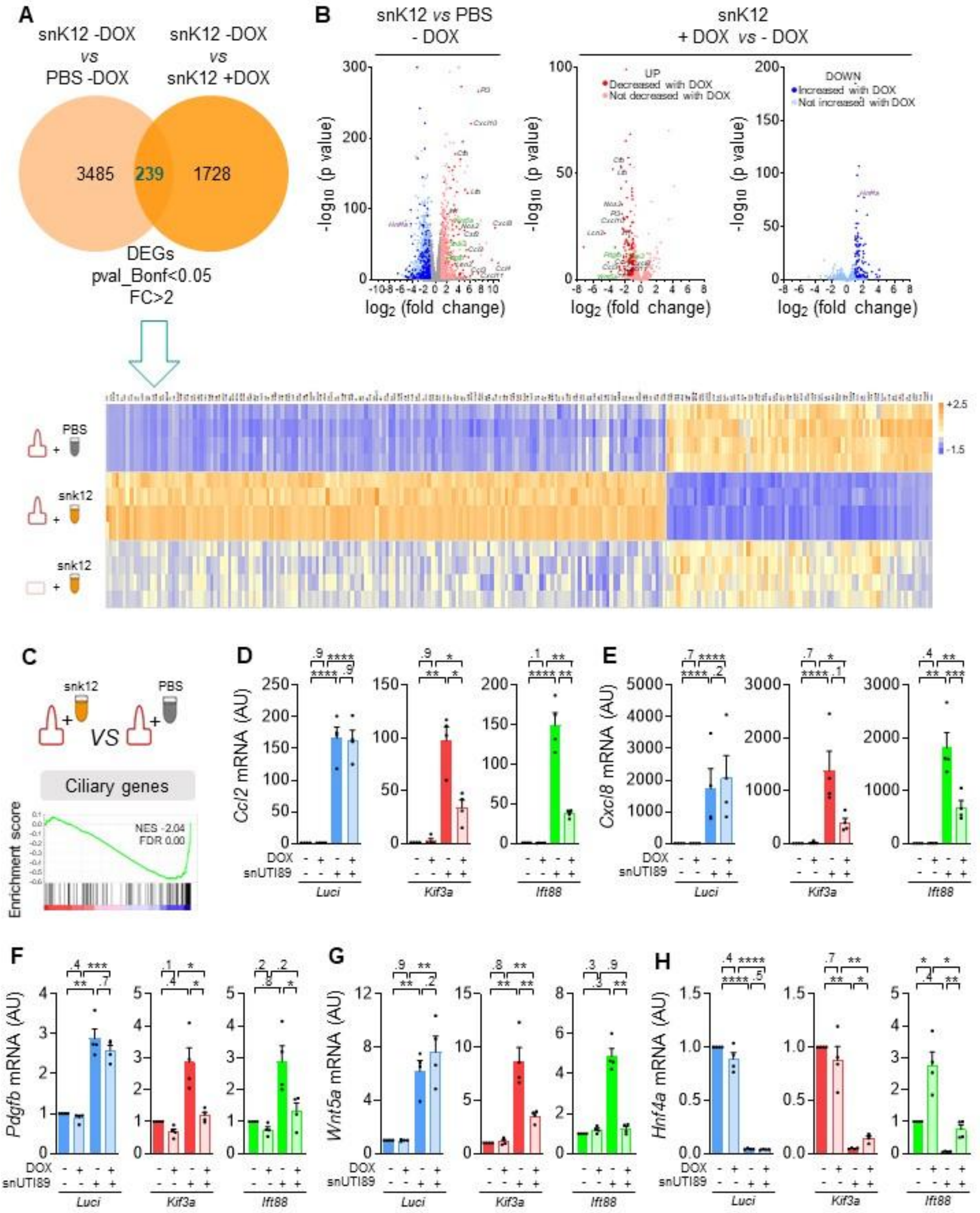

**Extended Data Figure 5.**

**(A)** Venn diagram representing the DEGs upon snK12 compare to vehicle (left circle), the impact of primary cilia loss upon doxycycline treatment on their expression (right circle) and the 239 genes rescued by primary cilia loss (intersection). These 239 DEGs are represented on the heatmap showing the computed Z-scores. **(B)** Volcano plots representation of the differentially expressed genes (DEGs) in 10 days cultured *Kif3a* i-shRNA MDCK cells treated for 6 hours with supernatant of heat killed *E. coli* (snK12) or vehicle stimulation (PBS, left side) and the impact of primary cilia loss upon doxycycline treatment on their expression (middle and right side). **(C)** Gene set enrichment analysis comparing selected for ciliary genes comparing the ciliated cells treated with *E. coli* extracts (snK12) or PBS. **(D-H)** qPCR quantifications of *Ccl2* (D), *Cxcl8* (E), *Pdgfb* (F), *Wnt5a* (G) and *Hnf4a* (H) mRNA in doxycycline (DOX) inducible *luciferase* (blue), *Kif3a* (red) or *Ift88* (green) shRNA MDCK cells treated for 6 hours with supernatant of heat killed UPEC (snUTI89) or vehicle (PBS). Each dot represents an independent experiment. Paired one-way RM ANOVA (log-normal) with Geisser Greenhouse correction followed by Tukey test: *$P < 0.05$, **$P < 0.01$, ***$P < 0.001$, ****$P<0.00001$. AU: arbitrary units.

Extended Data Fig. 6

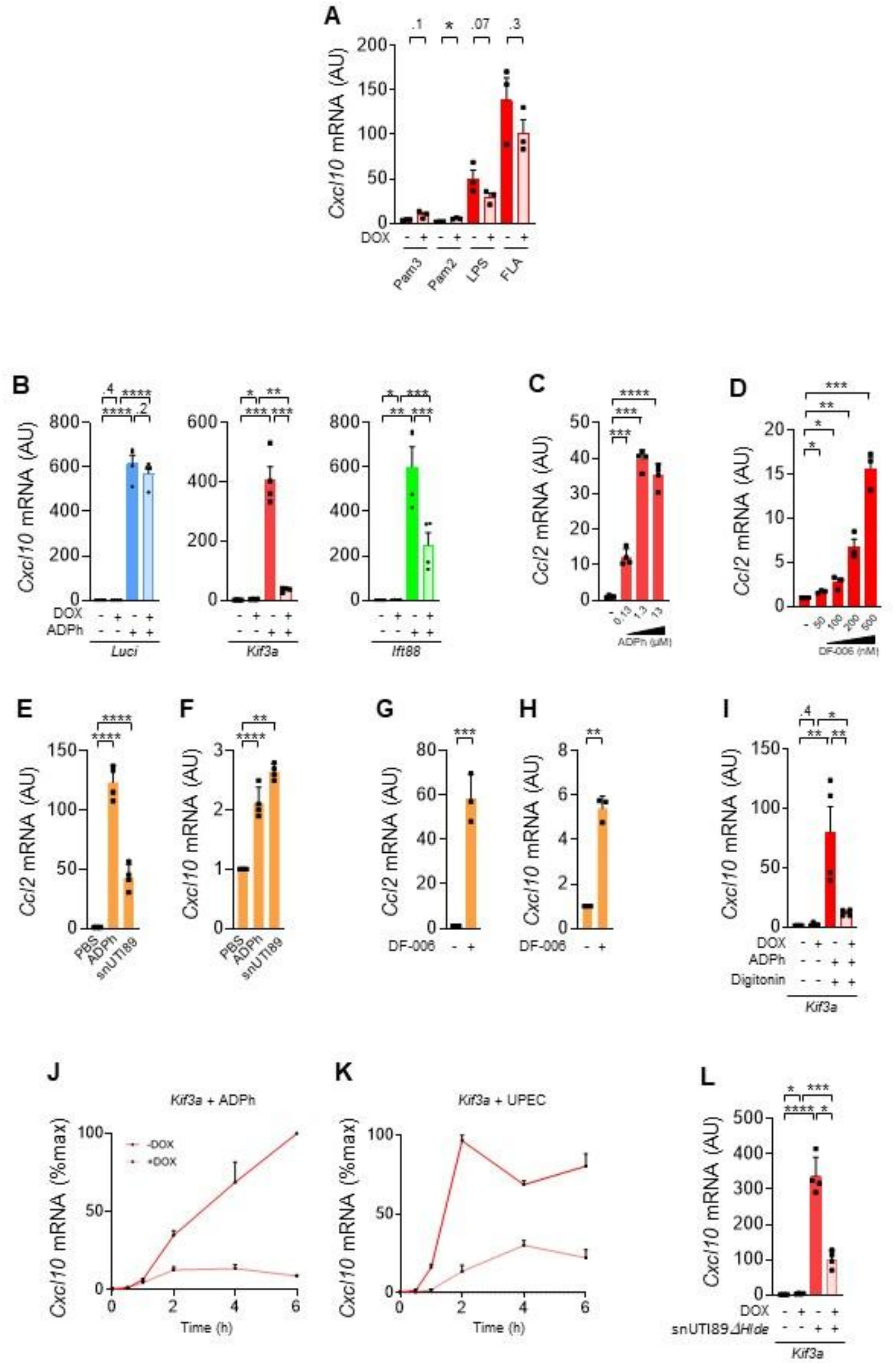

**Extended Data Figure 6.**

**(A)** qPCR quantification of *Cxcl10* mRNA in *Kif3a* i-shRNA MDCK cells treated for 6 hours with TLR-1/2 (Pam3), TRL-2/6 (Pam2), TLR-4 (Lipopolysaccharide, LPS) or TLR-5 agonist (Flagellin, FLA). **(B)** qPCR quantification of *Cxcl10* mRNA in *luciferase* (blue), *Kif3a* (red) or *Ift88* (green) i-shRNA MDCK cells treated for 6 hours with 13µM ADPh. **(C)** qPCR quantification of *Ccl2* mRNA in *Kif3a* i-shRNA MDCK cells treated for 6 hours with increasing doses of ADPh (0.13, 1.3 or 13µM). **(D)** qPCR quantification of *Ccl2* mRNA in *Kif3a* i-shRNA MDCK cells treated for 4 hours with increasing doses of DF-006 (50, 100, 200 or 500nM). **(E-F)** qPCR quantification of *Ccl2* (E) and *Cxcl10* (F) mRNA in mIMCD3 cells treated for 6 hours with ADPh (13µM) or supernatant of heat killed UPEC (snUTI89). **(G-H)** qPCR quantifications of *Ccl2* (G) and *Cxcl10* (H) mRNA in mIMCD3 cells treated for 4 hours with DF-006 (500nM). **(I)** qPCR quantification of *Cxcl10* mRNA in *Kif3a* i-shRNA MDCK cells treated for 30min with 13µM ADPh in presence of digitonin, and stopped 5 hours and a half later. **(J-K)** qPCR quantification of *Cxcl10* mRNA expression (expressed in percentage of the maximum value) depending on treatment duration (0min, 30min, 1h, 2h, 4h or 6h) after ADPh (J) and UPEC (K) stimulation of *Kif3a* i-shRNA MDCK cells. **(L)** qPCR quantification of *Cxcl10* mRNA in *Kif3a* i-shRNA MDCK cells treated for 6 hours with UPEC heat-killed supernatant lacking *Hlde* (snUTI89*ΔHlde*). **(A, C-H)** Ratio paired *t* test: *P<0.05, **P<0.01, ***P<0.001, ****P<0.00001. Each dot represents an experimental n. AU: arbitrary units. **(B, I, L)** Paired one-way RM ANOVA (log-normal) with Geisser Greenhouse correction followed by Tukey test: *P<0.05, **P<0.01, ***P<0.001, ****P<0.00001. Each dot represents an experimental n. AU: arbitrary units.

Extended Data Fig. 7

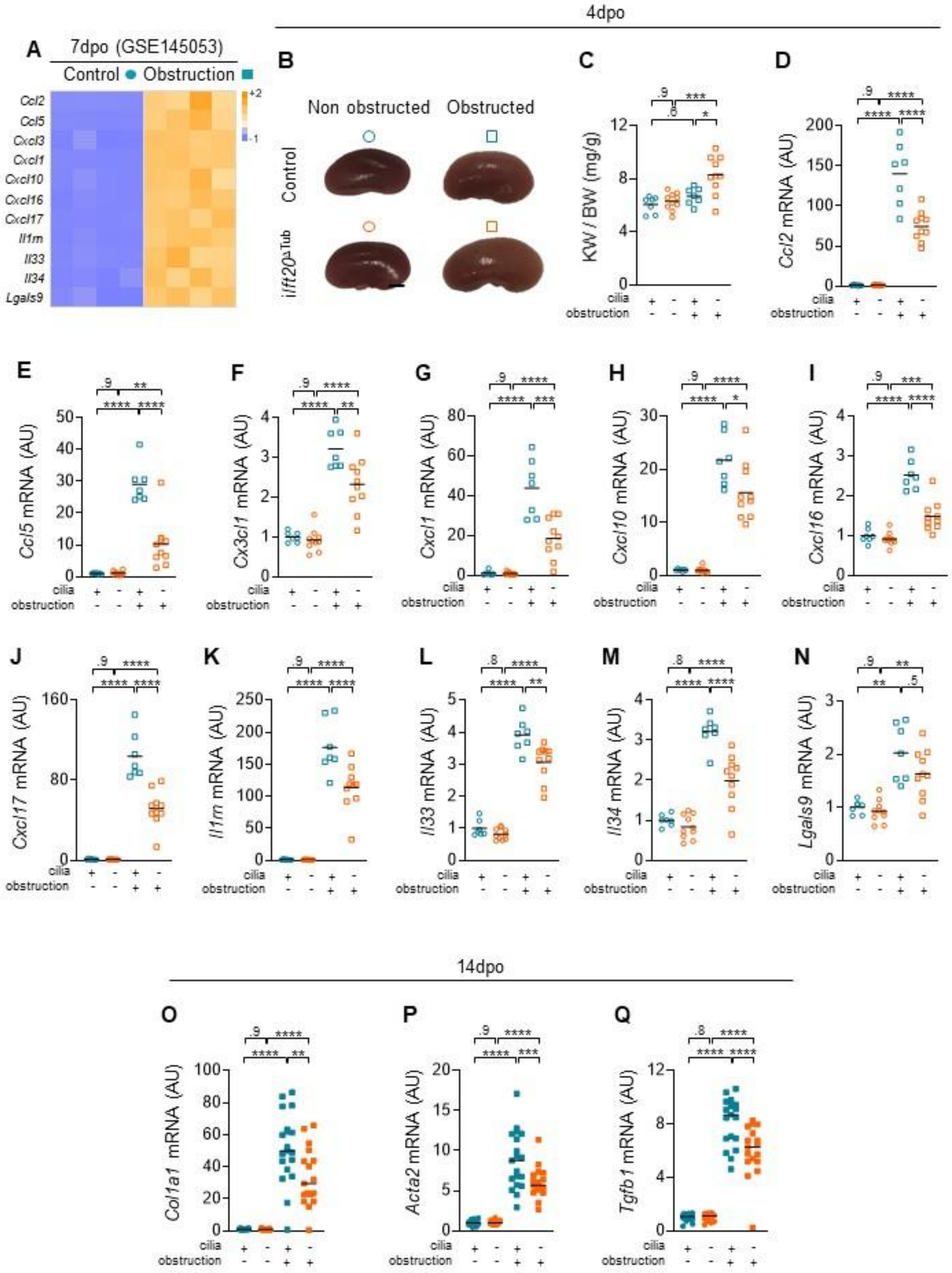

**Extended Data Figure 7.**

**(A)** Heatmap showing the Z-scores computed on published RNAseq normalized counts (GSE145053) of inflammatory transcripts in the obstructed kidneys from control seven days after surgery. **(B)** Representative pictures of the obstructed and non-obstructed kidneys from 10-week-old control and *Ift20*$^{\Delta Tub}$ mice, four days after surgery. Scale bar: 2mm. **(C)** Kidney weight (KW) to body weight (BW) ratio of the obstructed and non-obstructed kidneys from 10-week-old control and *Kif3a*$^{\Delta Tub}$ mice, four days after surgery. **(D-N)** qPCR quantification of *Ccl2* (D), *Ccl5* (E), *Cx3cl1* (F), *Cxcl1* (G), *Cxcl10* (H), *Cxcl16* (I), *Cxcl17* (J), *Il1rn* (K), *Il33* (L), *Il34* (M) and *Lgals9* (N) mRNA of the obstructed and non-obstructed kidneys from 10-week-old control and *Ift20*$^{\Delta Tub}$ mice, four days after surgery. **(O-Q)** qPCR quantification of *Col1a1* (O), *Acta2* (P) and *Tgfb1* (Q) mRNA of the obstructed and non-obstructed kidneys from 11-week-old control and *Ift20*$^{\Delta Tub}$ mice, fourteen days after surgery. **(B-Q)** Bars indicate mean. Each dot represents a mouse. Filled symbols represent females while empty symbols represent males. One-way ANOVA followed by Tukey test: *P<0.05, **P<0.01, ***P<0.001, ****P<0.00001. AU: arbitrary units.

Extended Data Fig. 8

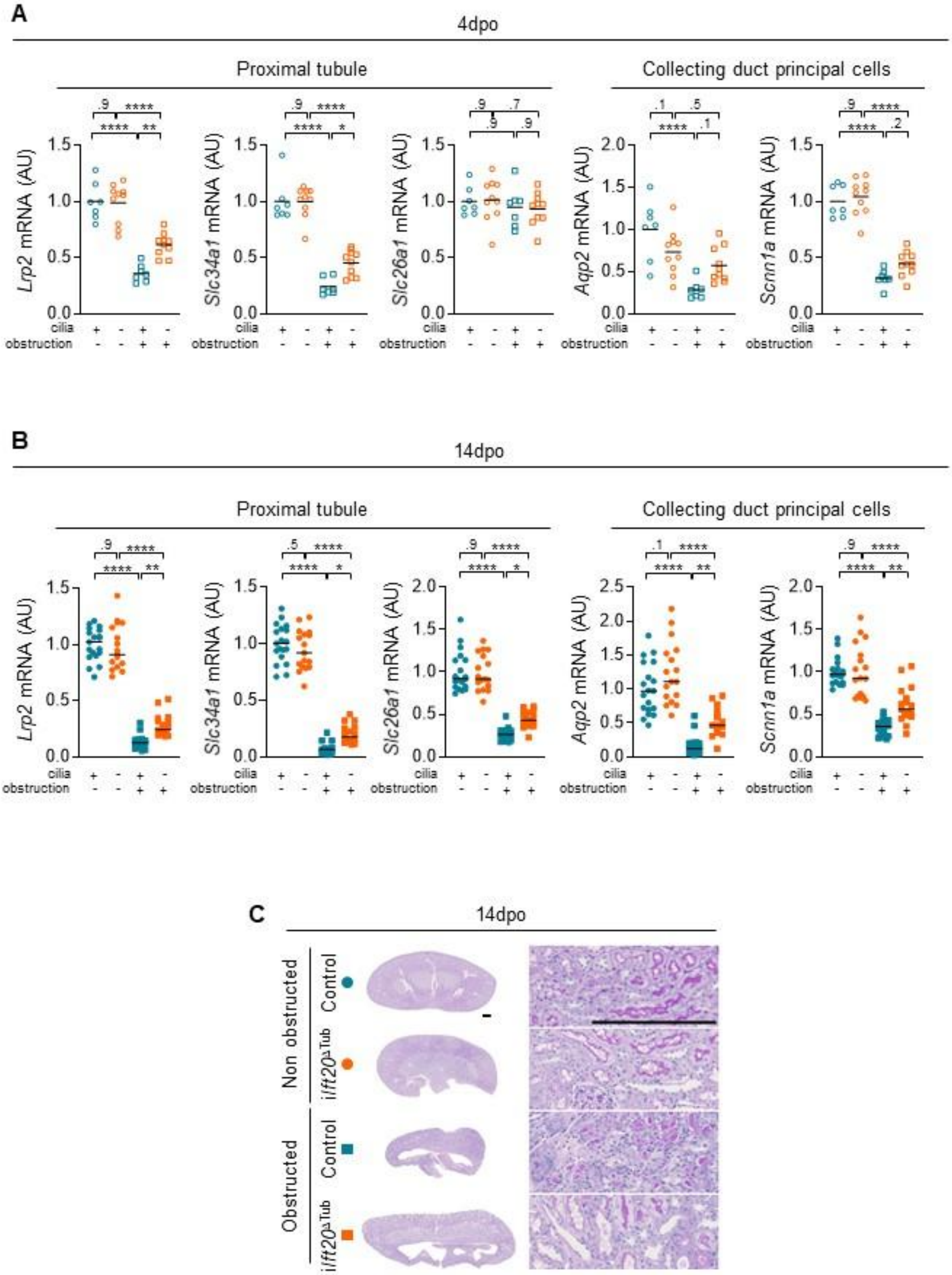

**Extended Data Figure 8.**

**(A-B)** qPCR quantification of proximal differentiation markers (*Lrp2*, *Slc34a1*, *Slc26a1*) and collecting duct principal markers (*Aqp2*, *Scnn1a*) mRNA of the obstructed and non-obstructed kidneys from control and *Ift20*$^{\Delta Tub}$ mice, four (B) and fourteen (C) days after surgery. Bars indicate mean. Each dot represents a mouse. Filled symbols represent females while empty symbols represent males. One-way ANOVA followed by Tukey test: *P<0.05, **P<0.01, ***P<0.001, ****P<0.00001. AU: arbitrary units. **(C)** Representative PAS staining of the obstructed and non-obstructed kidneys from 10-week-old control and *Ift20*$^{\Delta Tub}$ mice, fourteen days after surgery. Scale bar: 0.25mm.

Extended Data Fig. 9

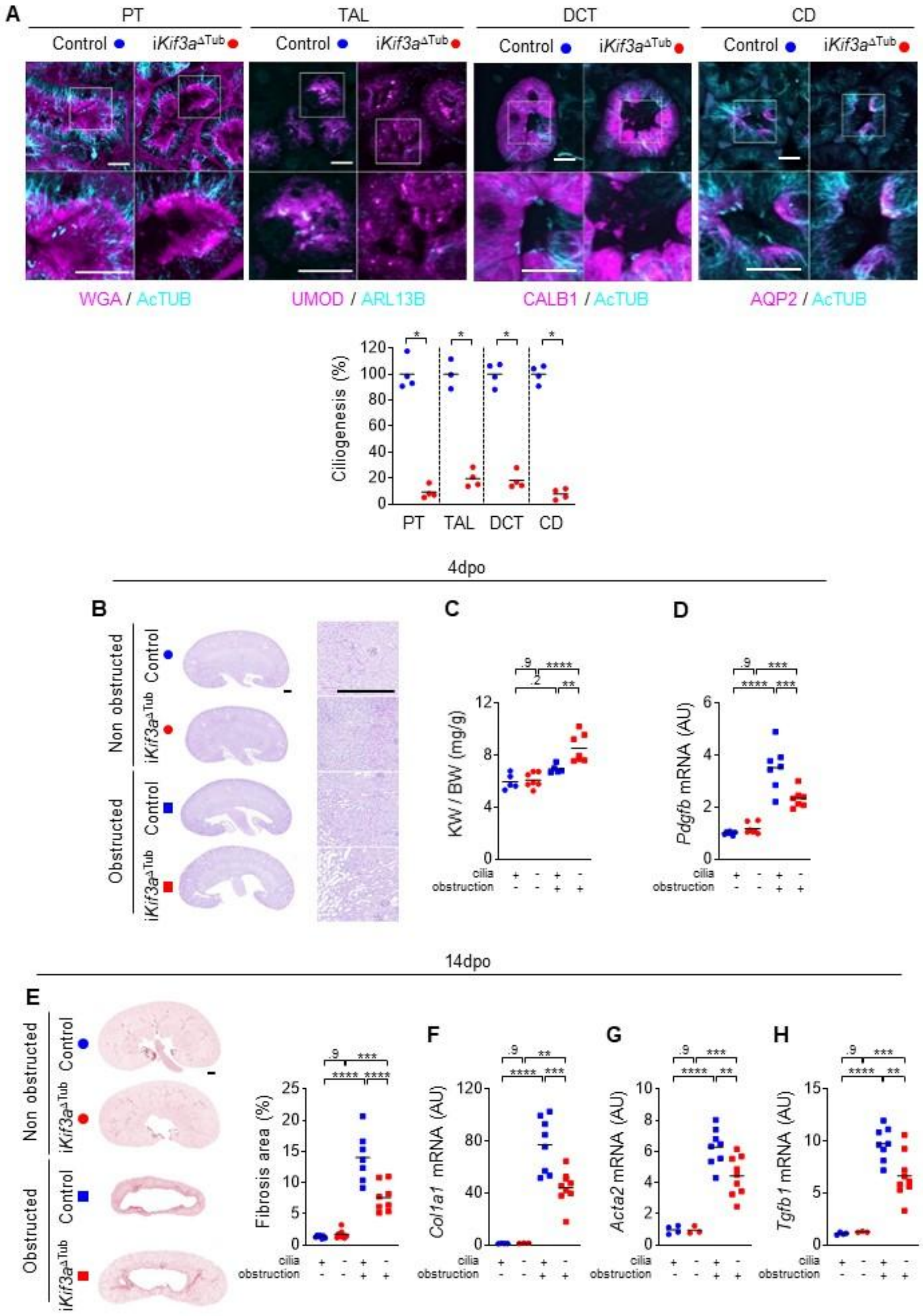

**Extended Data Figure 9.**

**(A)** Representative images and quantification of primary cilia (AcTUB+ or ARL13B+, blue) in proximal tubules (PT, with WGA+ brush border), thick ascending loop of Henle (TAL, UMOD+), distal convoluted tubules (DCT, CALB1+) and collecting duct principal cells (CD, AQP2+) of kidneys from 8-week-old control and *Kif3a*$^{\Delta\mathrm{Tub}}$ mice, 2 weeks after the completion of doxycycline treatment. Scale bar: 10µm. Bars indicate mean. Each dot represents a female mouse. Mann-Whitney t-test: *P<0.05. **(B)** Representative PAS staining of the obstructed and non-obstructed kidneys from 8-week-old control and *Kif3a*$^{\Delta\mathrm{Tub}}$ mice, four days after surgery. Scale bar: 0.5mm. **(C)** Kidney weight (KW) to body weight (BW) ratio of the same mice. **(D)** qPCR quantification of *Pdgfb* mRNA in the same mice. **(E)** Representative images and quantification of picrosirius stained fibrotic area on the obstructed and non-obstructed kidney sections from 10-week-old control and *Kif3a*$^{\Delta\mathrm{Tub}}$ mice, fourteen days after surgery. Scale bar: 0.5 mm. **(F-H)** qPCR quantification of *Col1a1* (F), *Acta2* (G) and *Tgfb1* (H) mRNA in kidneys from the same mice. **(C-D, F-H)** Bars indicate mean. Each dot represents a female mouse. One-way ANOVA followed by Tukey test: **P<0.01, ***P<0.001, ****P<0.00001. AU: arbitrary units.

Extended Data Fig. 10

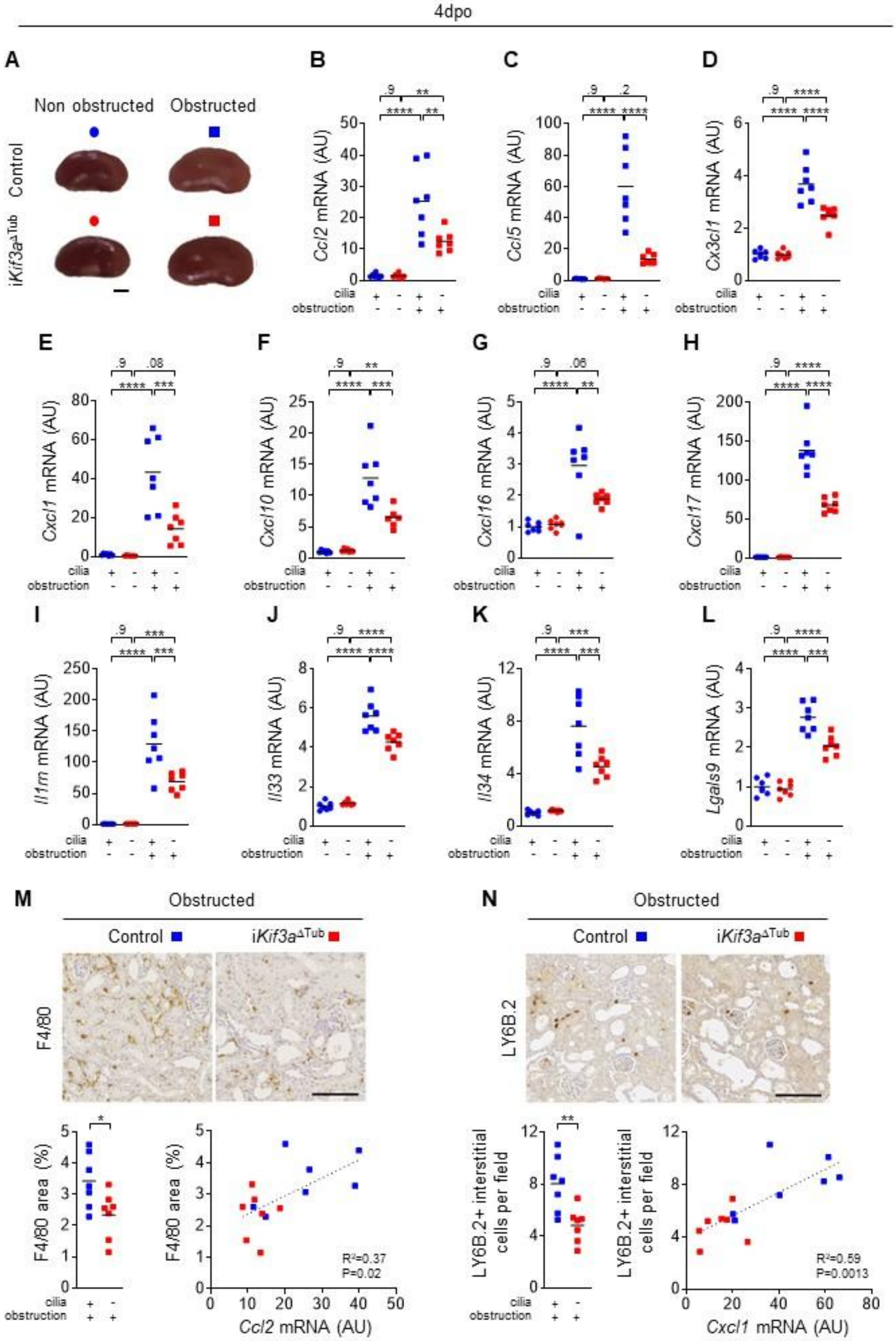

**Extended Data Figure 10.**

**(A)** Representative images of the obstructed and non-obstructed kidneys from 8-week-old control and *Kif3a*$^{\Delta Tub}$ mice, four days after surgery. Scale bar: 2mm. **(B-L)** qPCR quantification of *Ccl2* (B), *Ccl5* (C), *Cx3cl1* (D), *Cxcl1* (E), *Cxcl10* (F), *Cxcl16* (G), *Cxcl17* (H), *Il1rn* (I), *Il33* (J), *Il34* (K) and *Lgals9* (L) mRNA of the same mice. **(M)** Representative images and quantification of F4/80 (mononuclear phagocytes) staining and linear regression with *Ccl2* mRNA level in obstructed kidneys from 8-week-old control and *Kif3a*$^{\Delta Tub}$ mice, four days after surgery. Scale bars: 100 µm. **(E)** Representative images and quantification of LY6B.2 (neutrophils) staining and linear regression with *Cxcl1* mRNA level in obstructed kidneys from 8-week-old control and *Kif3a*$^{\Delta Tub}$ mice, four days after surgery. Scale bars: 100 µm. **(B-L)** Bars indicate mean. Each dot represents a female mouse. One-way ANOVA followed by Tukey test: **P<0.01, ***P<0.001, ****P<0.00001. AU: arbitrary units. **(M-N)** Bars indicate mean. Each dot represents a mouse. Mann-Whitney t-test : *P<0.05, **P<0.01.

Extended Data Fig. 11

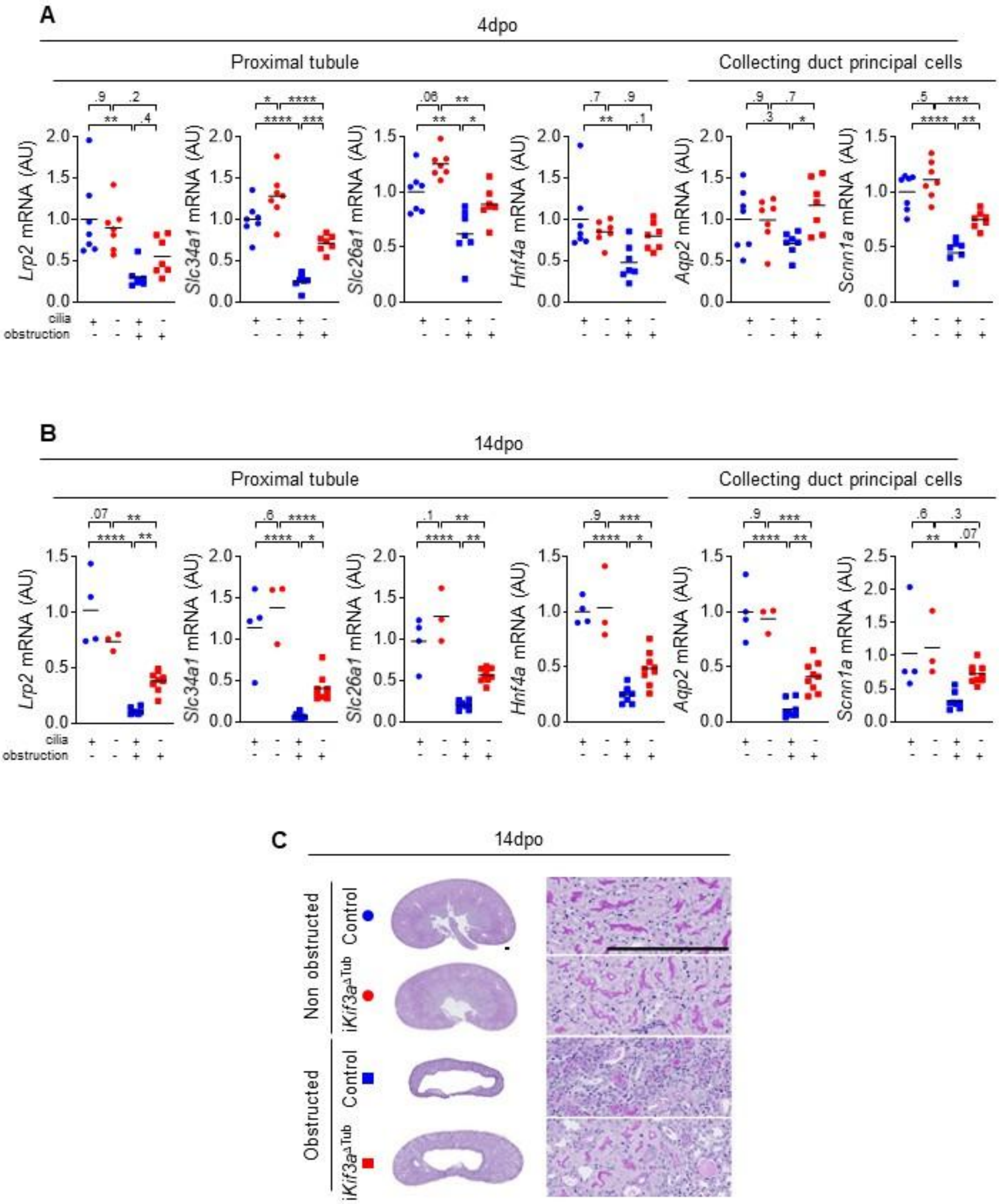

**Extended Data Figure 11.**

**(A-B)** qPCR quantification of proximal differentiation markers (*Lrp2*, *Slc34a1*, *Slc26a1*) and collecting duct principal markers (*Aqp2*, *Scnn1a*) mRNA of the obstructed and non-obstructed kidneys from control and *Kif3a*$^{\Delta Tub}$ mice, four (B) and fourteen (C) days after surgery. Bars indicate mean. Each dot represents a mouse. Filled symbols represent females. One-way ANOVA followed by Tukey test: *P<0.05, **P<0.01, ***P<0.001, ****P<0.00001. AU: arbitrary units. **(C)** Representative PAS staining of the obstructed and non-obstructed kidneys from 10-week-old control and *Kif3a*$^{\Delta Tub}$ mice, fourteen days after surgery. Scale bar: 0.25mm.

### Extended Data Fig. 12

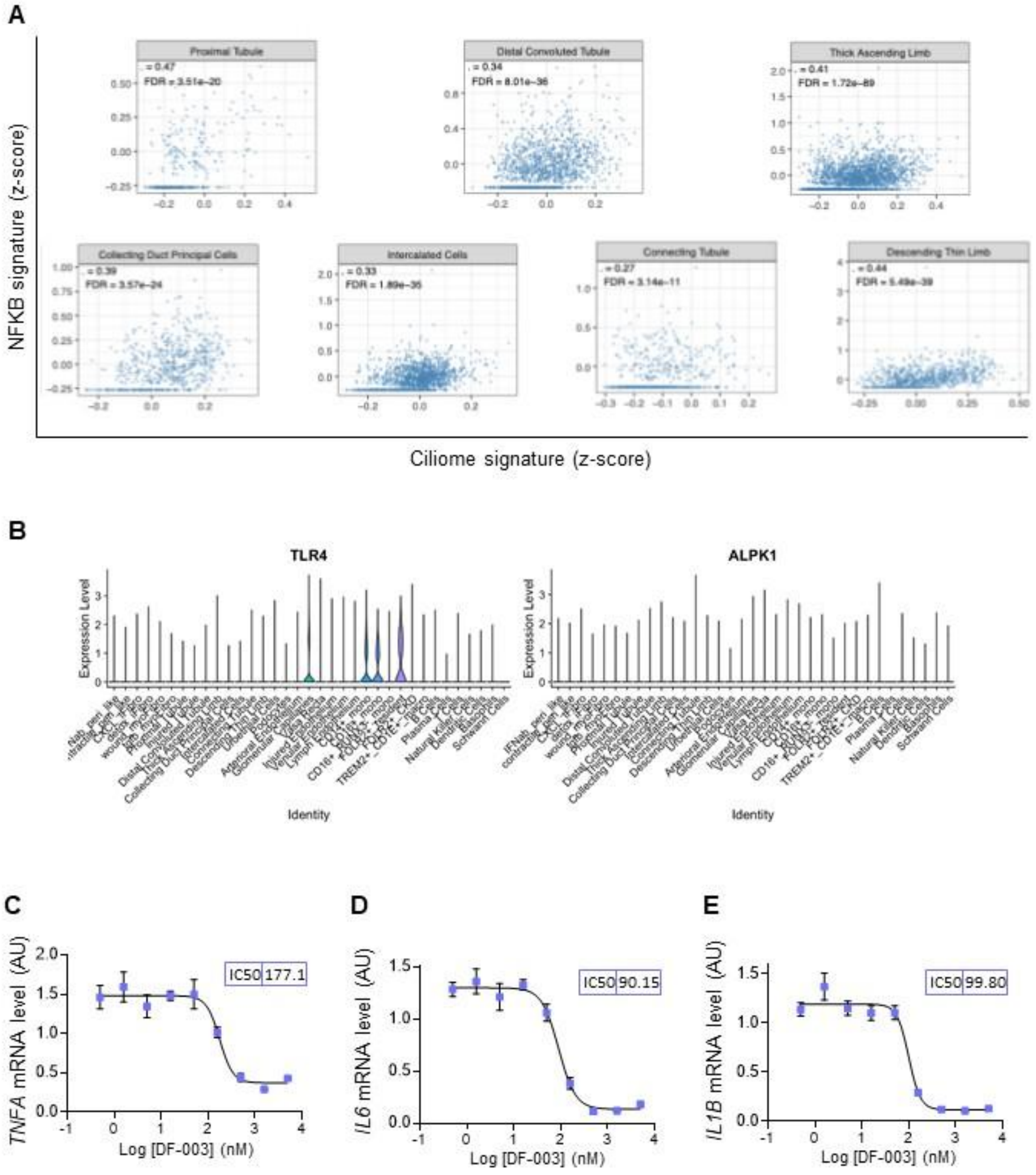

**Extended Data Figure 12.**

**(A)** Scatter plots of ciliome versus NF-κB pathway signature scores computed for each tubular cell type independently. Each dot represents a single cell. Spearman correlation coefficients (ρ) and Benjamini-Hochberg-adjusted p-values (FDR) are indicated for each population. All eight tubular cell types display significant positive correlations (ρ ranging from 0.27 in connecting tubule to 0.47 in proximal tubule; FDR < 0.05 for all). **(B)** *TLR4* and *ALPK1* expression were analyzed on single-cell RNA sequencing data from the Kramann human kidney atlas were analyzed. **(C-E)** qPCR quantification of *TNFa* (B), *IL6* (C) and *IL1B* (D) mRNA in human primary kidney epithelial cells treated for 4 h with DF-006 (100 nM) and pre-treated for 2 h with increasing doses of DF-003.
